# The RND efflux pump EefABC is highly conserved within lineages of *E. coli* commonly associated with infection

**DOI:** 10.1101/2024.08.01.606150

**Authors:** Hannah L. Pugh, Elizabeth M. Darby, Leah Burgess, Abigail L. Colclough, Asti-Rochelle Meosa John, Steven Dunn, Christopher Connor, Eoughin A. Perry, Alan McNally, Vassiliy N. Bavro, Jessica M. A. Blair

## Abstract

Tripartite resistance-nodulation-division (RND) efflux pumps confer multidrug resistance (MDR) in Gram-negative bacteria and are critical for many physiological functions including virulence and biofilm formation. The common laboratory strain of *E. coli,* K-12 MG1655 has six recognised RND transporters participating in tripartite pump formation (AcrB, AcrD, AcrF, CusA, MdtBC, and MdtF). However, by studying >20,000 *E. coli* genomes we show that *E. coli* belonging to phylogroups B2, D, E, F and G, which are commonly associated with infection, possess an additional, seventh RND transporter, EefB. It is found in a five gene operon, *eefRABCD,* which also encodes a TetR family transcription factor, a periplasmic adapter protein, an outer membrane factor and major facilitator superfamily pump. In contrast, *E. coli* from phylogroups A, B1 and C, generally containing environmental and commensal strains, do not encode the operon and instead encode an uncharacterised ORF, *ycjD*. In phylogroups where the *eefRABCD* operon is present it was very highly conserved. In fact, conservation levels were comparable to that of the major *E. coli* RND efflux system AcrAB-TolC, suggesting a critical biological function. Protein modelling shows that this pump is highly divergent from endogenous *E. coli* RND systems with unique structural features, while showing similarities to efflux systems found in *Pseudomonas aeruginosa*. However, unlike other major RND efflux systems, EefABC does not appear to transport antimicrobials and instead may be important for infection or survival in the host environment.

**Importance:** Efflux pumps are molecular machines that export molecules out of bacterial cells. The efflux pumps belonging to the RND family are particularly important as they export antibiotics out of Gram-negative bacterial cells, contributing to antibiotic resistance. The important human pathogen, *E. coli*, has been previously reported to have six RND pumps. However, we show that phylogroups of *E. coli* commonly associated with infection encode a seventh RND pump, EefABC which is highly conserved, suggesting an important biological function. While the function of EefABC in *E. coli* remains to be resolved, it does not seem to transport antimicrobial compounds. These findings are important because they reveal a new RND pump, potentially involved in virulence and survival in the host, that could represent a new therapeutic target. Additionally, it again shows that laboratory type strains of common bacterial pathogens are not representative of those that are infection causing.

## Introduction

*Escherichia coli* is a leading cause of invasive bacterial infections in humans, causing a range of diseases from urinary tract infections to haemorrhagic shock (Holmes et al., 2021). However, *E. coli* is also found in the wider environment and is a common commensal, colonising the gastrointestinal tract of both humans and animals. *E. coli* is a genetically diverse species that is divided into phylogroups (A, B1, B2, C, D, E, F and G) which are determined by genetic similarity (Denamur et al., 2021). Though virulence is not limited to specific phylogroups (Clermont et al., 2000, Gordon and Cowling, 2003), extra-intestinal infection is most commonly associated with phylogroups B2 and D (Denamur et al., 2021), while commensal or environmental lifestyles are mainly associated with phylogroups A and B1 (Tenaillon et al., 2010). In addition to phylogroups, *E. coli* isolates are also classified by sequence types (ST), determined by multi-locus sequence typing (MLST) (Maiden et al., 1998). Pandemic clonal STs such as ST131 are responsible for high incidence rates of extra-intestinal *E. coli* infections and the carriage of genes conferring multidrug resistance (Shaik et al., 2017, Li et al., 2023).

Gram-negative bacteria, such as *E. coli*, possess efflux pumps of the resistance-nodulation-division (RND) family which are best known for the export of antimicrobials and biocides (Chen et al., 2003, Sanchez et al., 1997, Nishino and Yamaguchi, 2001, Alav et al., 2021) with overexpression of these pumps conferring multidrug resistance (MDR) (Swick et al., 2011, Chetri et al., 2019). RND pumps form tripartite complexes with periplasmic adaptor proteins (PAP) and outer membrane factors (OMF) to span the inner membrane, periplasmic space, and the outer membrane (Alav et al., 2021). In addition to MDR, RND pumps have been implicated in a wide range of additional functions, including virulence (Nishino et al., 2006, Blair et al., 2015, Padilla et al., 2010), biofilm production (Baugh et al., 2014), decreased susceptibility to dyes, bile salts and fatty acids (Nishino and Yamaguchi, 2001, Mateus et al., 2021), export of polyamines and quorum sensing molecules (Chan et al., 2007), export of bacterial metabolites (Wang-Kan et al., 2021), copper and ion homeostasis (Grass and Rensing, 2001, Franke et al., 2001), and motility (Bazzini et al., 2011, Yang et al., 2011).

Many Gram-negative bacteria encode multiple RND efflux pumps, with differing substrate specificities which are often expressed only under specific environmental conditions. However, the number of RND pumps is highly variable between bacterial lineages. The human-restricted pathogen *Neisseria gonorrhoeae* possesses only a single RND pump (MtrCDE), while bacterial species that can colonise multiple habitats generally possess more. *Salmonella enterica* serovar Typhimurium has five RND pumps, whereas *E. coli* is generally reported to have six: AcrB, AcrD, AcrF, MdtBC, MdtF and CusA (Nishino and Yamaguchi, 2001). However, the number of RND pumps can also vary within a genus. For example, we have recently shown that species across the *Acinetobacter* genus possess between two and nine RND pumps, with species most-commonly associated with human infection tending to encode more RND efflux pumps (Darby et al., 2023). A study by Ma *et al*. found loss of MtrC function correlated with increased drug susceptibility in *N. gonorrhoeae* isolated from the cervical environment (Ma et al., 2020). Moreover, we recently demonstrated that AcrF in *E. coli* O157:H7 is non-functional due to a conserved insertion containing two stop codons (Pugh et al., 2023), showing that within a species not all efflux pumps present are always functional.

The expression of RND efflux pumps is controlled by a complex network of positive and negative regulators including repressors of the TetR family. We previously reported the presence of the TetR family regulator EefR in four of ten *E. coli* strains studied (Colclough et al., 2019). The *eefR* gene was encoded alongside genes annotated as *eefA* and *eefB* that are predicted to encode a PAP and RND pump, respectively. This RND pump was first reported in *Enterobacter aerogenes* (now reclassified as *Klebsiella aerogenes*), where *eefA, eefB* and *eefC* were found in a three gene operon without the regulator, *eefR* (Masi et al., 2005). In that study, the *eefABC* operon was not expressed under laboratory conditions due to transcriptional silencing by H-NS (Masi et al., 2005), however overexpression was found to confer resistance to erythromycin (Masi et al., 2006). The EefABC RND pump has also been reported in *Klebsiella pneumoniae,* where it has been linked to virulence, as deletion of *eefA* was found to reduce both colonisation of the gastrointestinal tract and tolerance to low pH (Coudeyras et al., 2008).

In *E. coli* the *eefABC* operon is absent from the widely studied strain K-12 but has been reported in a single highly drug resistant environmental isolate*, E. coli* SMS-3-5. In this isolate the efflux pump was part of a larger gene cluster which included the regulator *eefR* and a major facilitator superfamily (MFS) efflux pump *eefD* (Fricke et al., 2008). To date, the wider prevalence of the *eefRABCD* operon across the diversity of the *E. coli* population and its function in *E. coli* are both still unknown. Here we demonstrate that: 1) EefABC is exclusively found in clinically relevant phylogroups of *E. coli* and is highly conserved, 2) homology modelling reveals that the pump has several distinctive structural features compared to other RND efflux pumps, and 3) neither EefABC, nor EefD transport clinically relevant antimicrobials but can transport dyes.

## Materials and methods

### Prevalence and conservation of *eefRABCD* across *E. coli* and related species

The *eefB* gene from *E. coli* SMS-3-5 (CP000970.1) was aligned to *E. coli* genomes (taxid 562) within the NCBI RefSeq Genome Database.

A total of 20,013 *E. coli* genome assemblies from 38 STs (Achtman scheme) (Wirth et al., 2006) were downloaded from Enterobase (Zhou et al., 2020) for the determination of *eef* conservation across *E. coli*. Duplicates were removed based upon MASH (v 2.2.2) distances (Ondov et al., 2016). A custom blast database (BLAST v 2.10.0) (Camacho et al., 2009) was generated from the sequences of *eefR, eefA, eefB, eefC* and *eefD* of *E. coli* ATCC 25922 (CP009072.1) and used with ABRicate (v 0.9.8) (Seemann, 2017) to identify the presence and conservation of *eef* genes across the *E. coli* assemblies. Where an *eef* gene was found to be split across multiple contigs within an assembly, the assembly was removed from analysis as it was not possible to confirm from the assemblies alone whether this was due to a sequencing or assembly error, or the interruption of the gene.

The *eefRABCD* gene sequences from *E. coli* ATCC 25922 were also used to determine whether *Shigella* species encode the operon. Assemblies of *Shigella boydii* (*n* = 495)*, Shigella dysenteriae* (*n* = 497)*, Shigella flexneri* (*n* = 499), and *Shigella sonnei* (*n* = 500) were downloaded from Enterobase. As with *E. coli,* duplicates were identified using MASH and removed prior to running ABRicate. Assemblies containing *eef* genes split over multiple contigs were removed from the analysis as with *E. coli.* The *eefRABCD* genes from *E. coli* ATCC 25922 were also aligned to *Salmonella* (taxid 590)*, Photorhabdus* (taxid 29487)*, Yersinia* (taxid 629)*, Serratia* (taxid 613)*, Pseudomonas* (taxid 286), *Enterobacter* (taxid 547) and *Acinetobacter* (taxid 469) genomes to identify any homologs in related Gammaproteobacteria, however this was achieved using the NCBI RefSeq Genome Database.

### The phylogenetic context of *eefRABCD*

Five assemblies of each *E. coli* ST (excluding ST84 where *n =* 2) and *E. fergusonii* were chosen at random to generate a phylogenetic tree. Assemblies were annotated using Prokka (v 1.14.6) (Seemann, 2014) with subsequent GFF files used as input for Roary (v 3.13.0) (Page et al., 2015). The core gene alignment produced by Roary was used to construct a GTR-gamma tree with 100 bootstraps using RaXmL (v 8.2.12) (Stamatakis, 2014). Trees were visualised and annotated using iTOL (Letunic and Bork, 2019).

### Genomic context of *eefRABCD*

Due to limited availability of RefSeq genomes for some *E. coli* STs used in this work, seven were chosen at random. Reference sequences were downloaded from NCBI NC_004431.1 (ST73), NC_007946.1 (ST95), NZ_HG941718.1 (ST131), NC_002695.2 (ST11), NC_000913.3 (ST10), NC_011751.1 (ST69), and NZ_CP035350.1 (ST617). The location of the *eef* operon was identified in the ST73, ST95, ST131 and ST11 and downloaded along with the flanking 10,000 bp. The homologous regions in ST10, ST69 and ST617 were also identified and downloaded. Alignments of the genomic regions in all seven reference sequences was performed using EasyFig (v 2.2.2) (Sullivan et al., 2011).

Genomes of additional bacterial species; *E. albertii* (CP070290.2), *E. marmotae* (CP056165.1)*, K. aerogenes* (NZ_CP041925.1)*, K. pneumoniae* (NC_016845.1), *Enterobacter vonholyi* (VTUC01000001.1), *Enterobacter dykesii* (VTTY01000003.1), *Enterobacter wuhouensis* (SJOO01000006.1), *Enterobacter kobei* (KI973153.1), *Enterobacter chengduensis* (CP043318.1), *S. boydii* (CP026836.1)*, S. dysenteriae* (CP026774.1), *S. flexneri* (AE005674.2) and *S. sonnei* (CP055292.1) were downloaded from NCBI. The visualisation of the genomic context of the *eef* operon and homologs was achieved using EasyFig. Mapping of insertion sequences in *S. dysenteriae* was done using ISEScan (v 1.7.2.3) (Xie and Tang, 2017).

### Homology modelling and structural analysis

Multiple sequence alignments (MSA) were prepared using MAFFT and NJ/UPGMA phylogeny algorithms as implemented in MAFFT v.7 server (https://mafft.cbrc.jp/) (Katoh et al., 2019). Phylo.io was used for phylogenetic guide tree visualisation (Robinson et al., 2016). Structural annotations of the MSA sequences were done with Espript 3 (http://espript.ibcp.fr) (Robert and Gouet, 2014).

For homology modelling, I-TASSER (Yang et al., 2015) was used in manual mode with assignment of templates and structural alignment, supplemented by SWISS-MODEL (Waterhouse et al., 2018). The following structural templates have been used for the specific protein modelling. EefA modelling: MexA (Uniprot P52477) 2V4D.pdb (Symmons et al., 2009); AcrA (Uniprot P0AE06) 5V5S.pdb (Wang et al., 2017). AcrA was used as a template due to the smaller gaps in the alignment and better quality of available full-length template. EefB modelling: MexB (Uniprot P52002) 3W9I.pdb (Nakashima et al., 2013); AcrB (Uniprot P31224) 2GIF.pdb (Seeger et al., 2006). EefC modelling: TolC (Uniprot P02930) 1EK9.pdb (Koronakis et al., 2000); OprM (Uniprot Q51487) 4Y1K.pdb (Monlezun et al., 2015); OprJ (Uniprot Q51397) 5AZS.pdb (Yonehara et al., 2016); OprN (Uniprot Q9I0Y7) 5AZO.pdb, 5AZP.pdb (Yonehara et al., 2016); 5IUY.pdb (Ntsogo Enguéné et al., 2017). EefD modelling: EmrD (Uniprot P31442) 2GFP.pdb (Yin et al., 2006); MdfA (Uniprot P0AEY8) 4ZP0.pdb (Heng et al., 2015).

For the models of the protein oligomers and the complete EefABC tripartite pump, rigid-body structural docking of the homology models was used guided by the available cryo-EM structure (5O66.pdb; (Wang et al., 2017)), the results of which were cross-validated manually and obvious steric clashes removed using Coot (Emsley et al., 2010). Additional structural analysis and visualisation was performed with Pymol (PyMOL Molecular Graphics System, Version 1.71 Schrödinger, LLC).

### Strains, plasmids and culture conditions

All strains used in this work are listed in Table S1. Strains were grown in lysogeny broth (LB, Merck) at 37°C with aeration unless stated otherwise.

### Cloning of *eefABCD*

The *eefABC* operon was amplified from the chromosome of *E. coli* ATCC 25922 (NCTC 12241) using Q5 polymerase (New England Biolabs) and primers which incorporated the *Nde*I and *Xho*I restriction sites (Table S2). The amplicon was cloned into both pET21a (ampicillin resistant) and pET24a (kanamycin resistant) plasmids (Invitrogen) which are identical aside from their resistance cassette. No IPTG induction was used in this work.

The *eefD* gene was amplified from the *E. coli* ATCC 25922 chromosome using Q5 (New England Biolabs) and primers that incorporated the *ApaLI* and *PstI* restriction sites (Table S2). The amplicon was then cloned into pACYC177 (ATCC).

### Deletion of *eefABC* and *eefD* in *E. coli* ATCC 25922

Deletion of the genes encoding the RND system of the *eefRABCD* operon was achieved by homologous recombination (Kim et al., 2014). However, due to the size of the *eefABC* operon, first *eefB* was interrupted, followed by the remaining *eefA* and *eefC* genes. An *eefD* knockout was generated independently using the same method. All primers are listed in Table S2.

### Determination of minimum inhibition concentration (MIC) of antimicrobials, metals, and dyes

Bacterial susceptibility to a range of antimicrobials and dyes was determined using the agar doubling dilution method described by the Clinical and Laboratory Standards Institute (CLSI, 2020). *E. coli* ATCC 25922 was used as a control to confirm antimicrobial efficacy in line with EUCAST guidelines (EUCAST, 2021). For the susceptibility to metals, bile salts and polyamines, a broth microdilution method was used (EUCAST, 2021).

### Accumulation and efflux of ethidium bromide

Ethidium bromide (EtBr) accumulation was measured as previously described (Smith and Blair, 2014). Briefly, EtBr was added to cells and the increase in fluorescence was measured over time.

Efflux activity was also assessed as previously described (Smith and Blair, 2014). Here, cells were incubated in the presence of EtBr and carbonyl cyanine m-chlorophenyl hydrazone (CCCP) until fluorescence saturation was reached. Re-energisation was achieved with glucose and the rate of reduction in fluorescence was measured.

## Results

### The *eefRABCD* operon is present in *E. coli* phylogroups associated with infection

To first understand how widespread the *eefRABCD* operon is across *E. coli,* the NCBI RefSeq Genome Database was utilised. The *eefB* gene from *E. coli* SMS-3-5 was aligned to genomes belonging to *E. coli* (taxid 562) using NCBI nucleotide blast. The top 100 matches in the *eefB* alignment had 100% sequence coverage and ≧97.7% sequence identity (data not shown), suggesting that the *eefB* gene was present more widely across *E. coli*. One of the strains found to possess *eefB* was the well characterised strain ATCC 25922. The *eef* operon in both *E. coli* SMS-3-5 and ATCC 25922 were found to be very similar and as a result, the operon from ATCC 25922 was used as the reference in future work.

Next we looked at the distribution of *eefRABCD* across *E. coli*. 20,013 assemblies were downloaded from the Enterobase database representing 38 STs. Using MASH distances, 766 assemblies were identified as duplicates and removed from the data set. The presence of *eefR, eefA, eefB, eefC* and *eefD* in the remaining 19,247 assemblies was determined using ABRicate (Table S3). Interestingly there was a clear divide between phylogroups where the *eefRABCD* operon was identified and those where it was completely absent (Fig. 1). The operon was not found in ST groups belonging to phylogroups A, B1 and C, which are more traditionally classified as environmental isolates. However, the *eefRABCD* operon was present in all ST groups of phylogroups B2, F and G, which are strongly associated with human infection and MDR. Notably, a single ST within phylogroups D and E lacked the operon, ST69 and ST182 respectively. Despite these two exceptions, a distinct divide between the A-B1-C and B2-D-E-F-G clades was identified, implying a clear evolutionary relationship.

**Figure 1.**
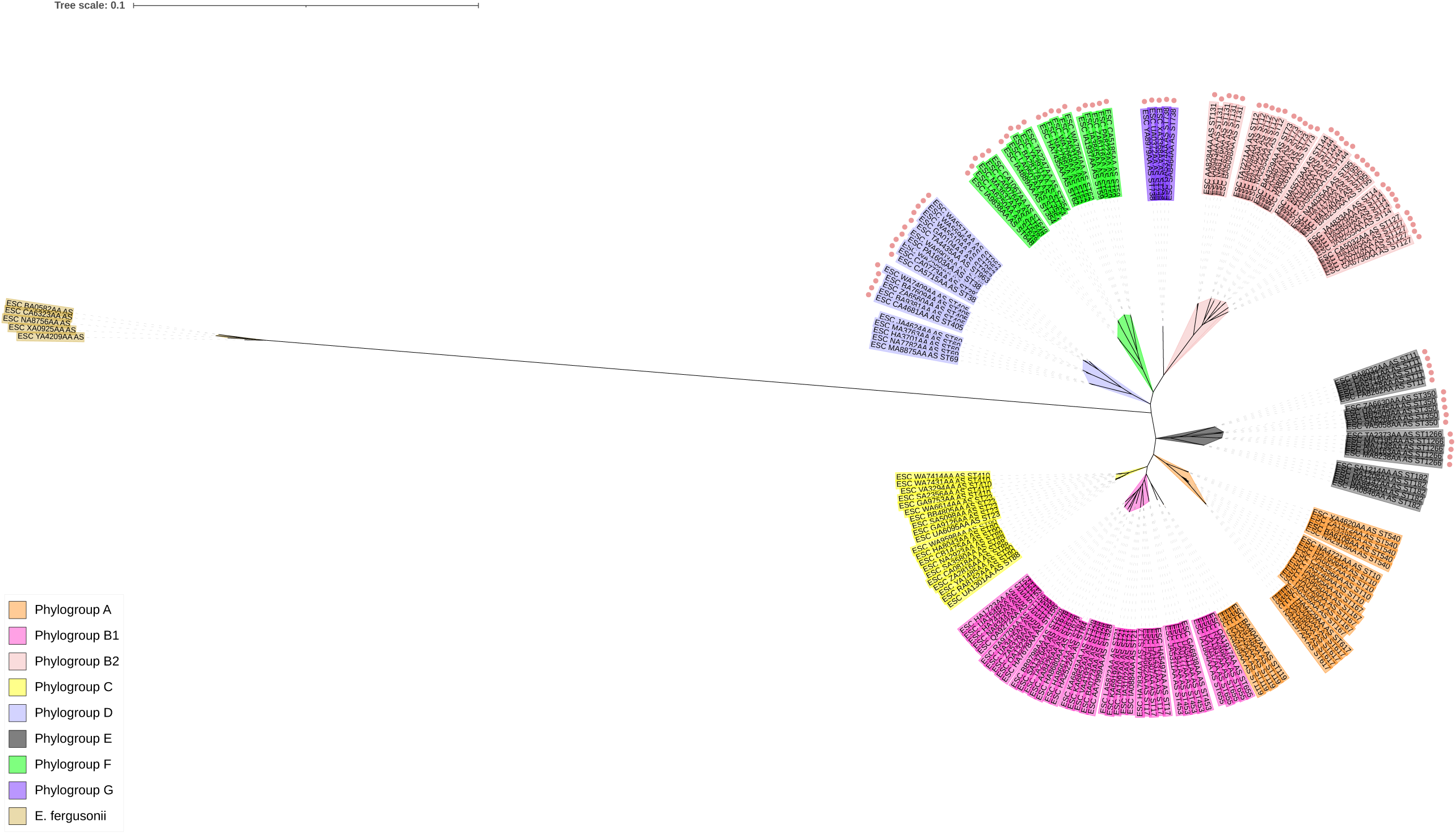
Phylogenetic structure of the assemblies used in this analysis and the distribution of *eefRABCD*. The tree was created using five assemblies per ST (exception ST84 *n=2*). Assemblies were chosen randomly. The tree was rooted using five *E. fergusonii* assemblies. The leaves are annotated with the ST group of the assembly and coloured coded by phylogroup, colours used to highlight phylogroups are shown in the legend on the bottom left corner. STs positive for the *eefRABCD* operon are marked with ●. Tree was generated using RaXmL and visualised on iTol.

### The *eefRABCD* operon is highly conserved across phylogroups B2, D, E, F and G

In *E. coli* assemblies positive for an *eefRABCD* gene, the entire operon was always present, with gene sequences highly conserved. Using *E. coli* ATCC 25922 as a reference, gene coverage for each component of the *eef* operon was always greater that 98% (Table S6). The nucleotide percentage identity varied between genes and across phylogroups though despite subtle variations, *eefR, eefA, eefB* and *eefD* averaged >99% overall. Interestingly, *eefC,* the gene coding for the OMF, was marginally more variable than the four other genes within the operon (Table 1).

**Table 1.**
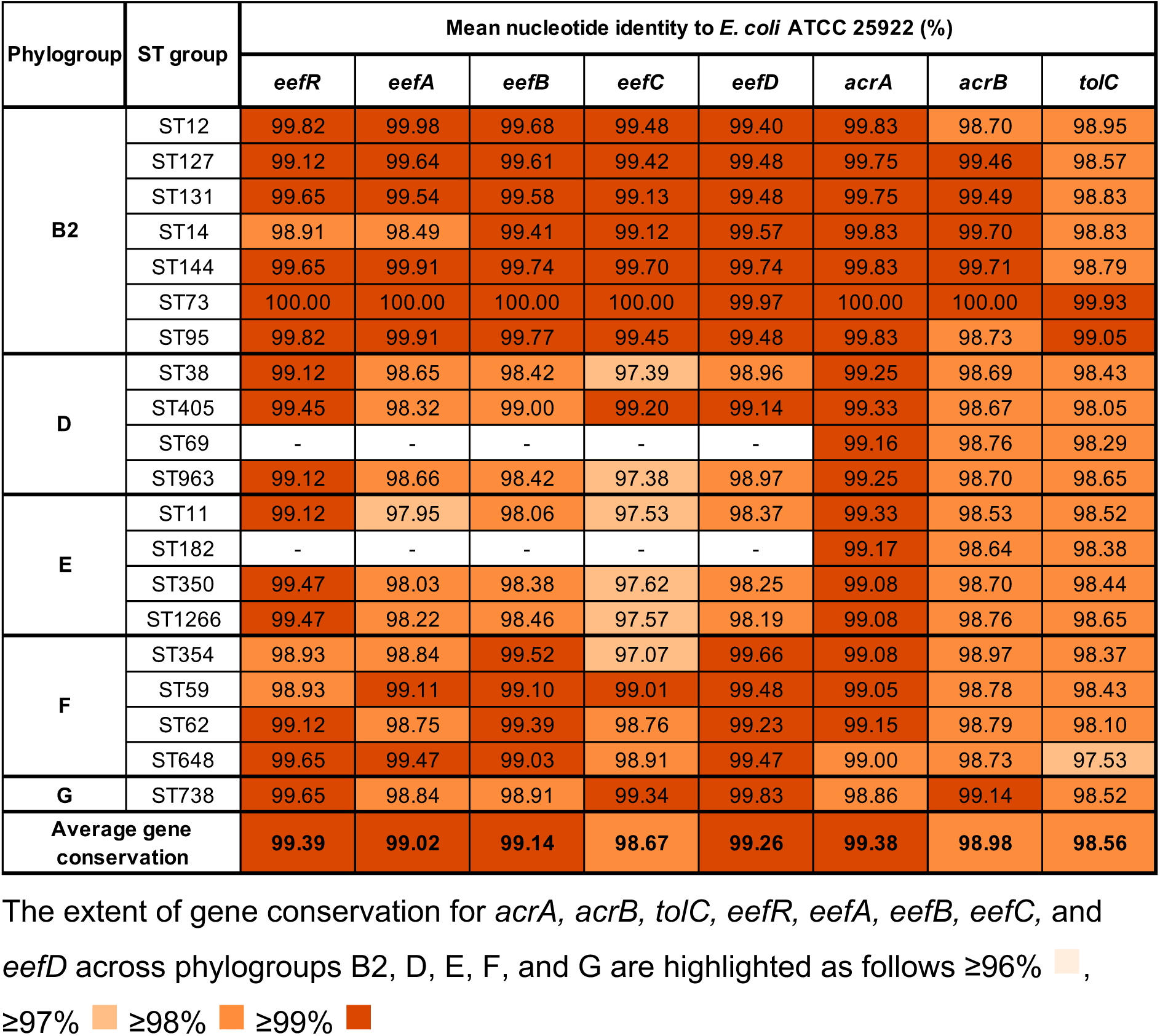
Conservation of *acrAB-tolC* and *eefRABCD* across *E. coli* phylogroups.

| Phylogroup | ST group | Mean nucleotide identity to *E. coli* ATCC 25922 (%) |  |  |  |  |  |  |  |
| --- | --- | --- | --- | --- | --- | --- | --- | --- | --- |
|  |  | *eefR* | *eefA* | *eefB* | *eefC* | *eefD* | *acrA* | *acrB* | *tolC* |
| **B2** | ST12 | 99.82 | 99.98 | 99.68 | 99.48 | 99.40 | 99.83 | 98.70 | 98.95 |
|  | ST127 | 99.12 | 99.64 | 99.61 | 99.42 | 99.48 | 99.75 | 99.46 | 98.57 |
|  | ST131 | 99.65 | 99.54 | 99.58 | 99.13 | 99.48 | 99.75 | 99.49 | 98.83 |
|  | ST14 | 98.91 | 98.49 | 99.41 | 99.12 | 99.57 | 99.83 | 99.70 | 98.83 |
|  | ST144 | 99.65 | 99.91 | 99.74 | 99.70 | 99.74 | 99.83 | 99.71 | 98.79 |
|  | ST73 | 100.00 | 100.00 | 100.00 | 100.00 | 99.97 | 100.00 | 100.00 | 99.93 |
|  | ST95 | 99.82 | 99.91 | 99.77 | 99.45 | 99.48 | 99.83 | 98.73 | 99.05 |
| **D** | ST38 | 99.12 | 98.65 | 98.42 | 97.39 | 98.96 | 99.25 | 98.69 | 98.43 |
|  | ST405 | 99.45 | 98.32 | 99.00 | 99.20 | 99.14 | 99.33 | 98.67 | 98.05 |
|  | ST69 | - | - | - | - | - | 99.16 | 98.76 | 98.29 |
|  | ST963 | 99.12 | 98.66 | 98.42 | 97.38 | 98.97 | 99.25 | 98.70 | 98.65 |
| **E** | ST11 | 99.12 | 97.95 | 98.06 | 97.53 | 98.37 | 99.33 | 98.53 | 98.52 |
|  | ST182 | - | - | - | - | - | 99.17 | 98.64 | 98.38 |
|  | ST350 | 99.47 | 98.03 | 98.38 | 97.62 | 98.25 | 99.08 | 98.70 | 98.44 |
|  | ST1266 | 99.47 | 98.22 | 98.46 | 97.57 | 98.19 | 99.08 | 98.76 | 98.65 |
| **F** | ST354 | 98.93 | 98.84 | 99.52 | 97.07 | 99.66 | 99.08 | 98.97 | 98.37 |
|  | ST59 | 98.93 | 99.11 | 99.10 | 99.01 | 99.48 | 99.05 | 98.78 | 98.43 |
|  | ST62 | 99.12 | 98.75 | 99.39 | 98.76 | 99.23 | 99.15 | 98.79 | 98.10 |
|  | ST648 | 99.65 | 99.47 | 99.03 | 98.91 | 99.47 | 99.00 | 98.73 | 97.53 |
| **G** | ST738 | 99.65 | 98.84 | 98.91 | 99.34 | 99.83 | 98.86 | 99.14 | 98.52 |
| **Average gene conservation** |  | **99.39** | **99.02** | **99.14** | **98.67** | **99.26** | **99.38** | **98.98** | **98.56** |

The extent of gene conservation for *acrA*, *acrB*, *tolC*, *eefR*, *eefA*, *eefB*, *eefC*, and *eefD* across phylogroups B2, D, E, F, and G are highlighted as follows $\geq 96\%$ , $\geq 97\%$ $\geq 98\%$ $\geq 99\%$

To contextualise the extent of *eefRABCD* conservation, the conservation of *acrA, acrB* and *tolC*, which encode the critically important RND efflux pump AcrAB-TolC in *E. coli*, was determined. In general, the nucleotide identity of the *eef* operon was conserved to a similar level as *acrAB-tolC*, all genes (excluding *eefC* in ST354) were >97% identical to those in *E. coli* ATCC 25922 (Table 1). However, conservation of both the *acrAB-tolC* and *eefABC* RND systems varied slightly at both the ST and phylogroup level. This suggests a strong selection pressure on the operon, indicating an important biological function.

Phylogroup B2 contains many clinically significant *E. coli* STs including the MDR ST131. Across phylogroup B2, *eefB was* generally more conserved than *acrB* with the *eef* genes having higher homology to the reference than the *acrAB-tolC* genes. However, in phylogroups D, E and F the opposite was seen as the *acrAB-tolC* genes were more conserved than the *eefRABCD* genes. Though only one phylogroup G ST was used in the analysis, in ST738 *eefR, eefC* and *eefD* had the highest homology to the reference, with *acrA, acrB, eefA* and *eefB* all equally conserved.

### Phylogroups A, B1 and C have highly conserved *sapF*-*fabI* intergenic region in place of *eefRABCD*

While *eefRABCD* was found to be highly conserved across phylogroups B2, D, E, F and G, it was completely absent from phylogroups A, B1 and C. To explore this further RefSeq genomes of strains with and without the operon were downloaded from NCBI and aligned using EasyFig (Fig. 2). In assemblies encoding the *eef* operon, it was always found at the same genomic location; that is between the essential gene *fabI* and the non-essential gene *sapF*. FabI is an enoyl-[acyl-carrier-protein] reductase that is involved in fatty acid production (Bergler et al., 1994), while SapF is a putrescine export protein belonging to the *sapBCDF* system (Sugiyama et al., 2016). Interestingly, in assemblies where the *eefRABCD* operon was absent, a hypothetical gene, *ycjD,* was annotated as present in the same genomic location. The *ycjD* open reading frame (ORF) was 354 nucleotides long and ran in the opposite orientation to *eefRABCD*.

**Figure 2.**
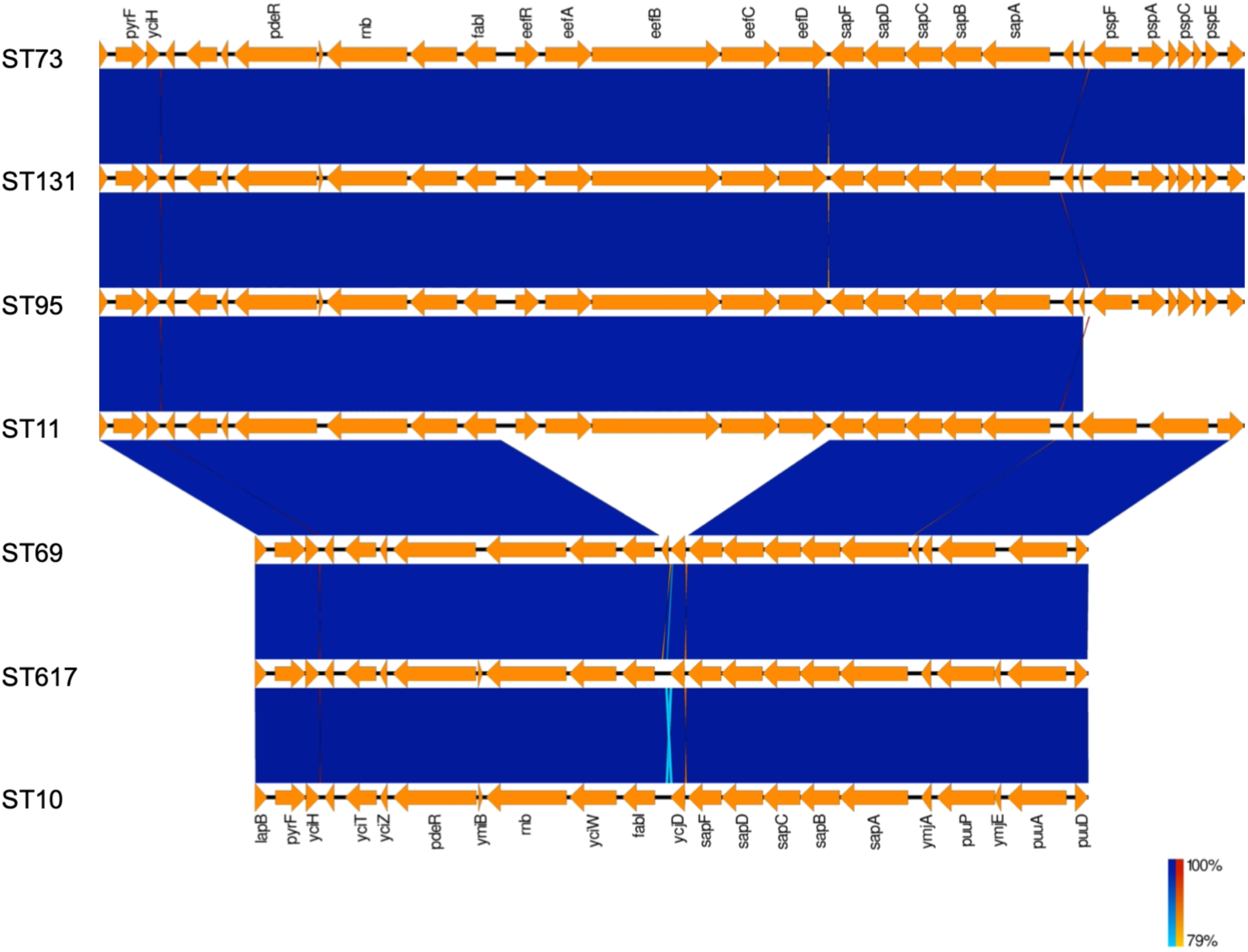
The genomic region of strains with and without *eefRABCD*. EasyFig alignment of the *eef* region in ST73 (B2), ST131 (B2), ST95 (B2), ST11 (E), ST69 (D), ST617 (A), ST10 (A). The *eefRABCD* operon was consistently identified between *fabI* and *sapF*.

A larger alignment of the 97 assemblies from phylogroups A, B1 and C (and ST69 and ST182) used to construct the phylogenetic tree demonstrated that the *ycjD* hypothetical gene is highly conserved across phylogroups A, B1 and C (Supplementary Data file). The alignment of the *sapF-ycjD-fabI* region also demonstrated that ST69 and ST182 possess an intergenic region highly homologous to the STs belonging to phylogroups A, B1 and C, though subtle differences were present (Fig. 3). The open reading frame annotated as *ycjD* in K-12, is 12 nucleotides longer in both ST69 and ST182 assemblies (Fig. 3). Taken together, these findings suggest that *eef* has been lost from *E. coli* on at least two independent instances.

**Figure 3.**
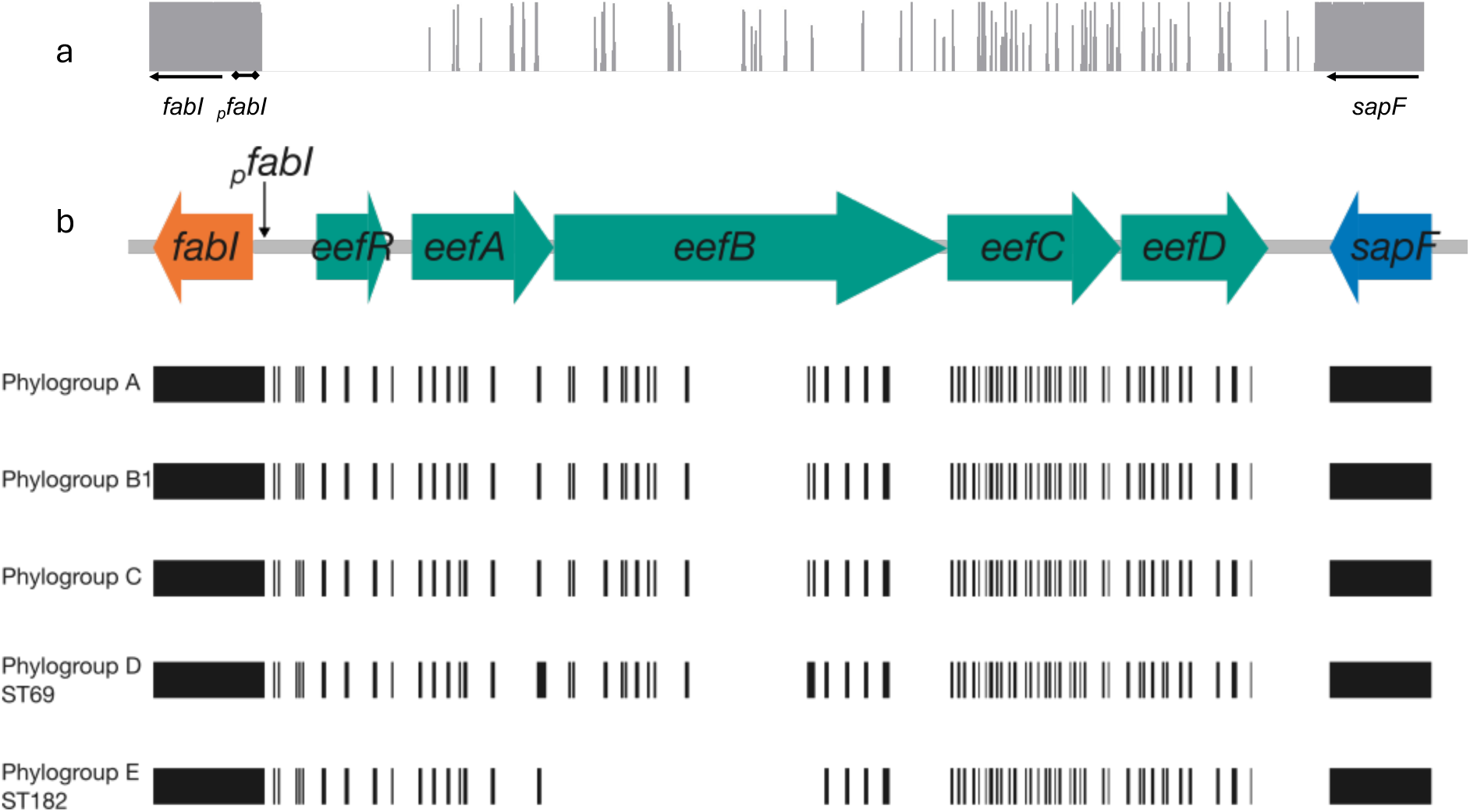
Diagrammatic representation of *eefRABCD* homology in strains that lack the operon. In *E. coli* ST positive for *eefRABCD*, the operon was consistently located between *fabI* and *sapF*. To determine whether *E. coli* assemblies that lacked the operon had conserved regions of the *eef* operon, the equivalent *fabI-sapF* intergenic region of assemblies negative for *eefRABCD* were aligned with *fabI*-*eefRABCD*-*sapF* from *E. coli* ATCC 25922. **A) Consensus identity of the *fabI-sapF* intergenic region in *E. coli* that lack the *eef* operon.** Between all STs, and despite phylogroup, the *fabI-sapF* intergenic regions aligned to the *eefRABCD* operon in a highly conserved manner. **B) Cartoon representation of operon fragment conservation in *E. coli* ST that lack the *eef* operon.** Closer inspection of the alignment (Supplementary file 1) found that phylogroups A, B1 and C had almost identical *fabI-sapF* intergenic regions, while ST69 and ST189 had marginally different homology patterns. Taken together, these data suggest the operon may have been lost in up to three independent events.

### Distribution of *eefRABCD* in Gram-negative bacteria

To see if the *eefRABCD* operon was present more widely across *Escherichia* species, the ATCC 25922 *eef* operon was aligned to sequences from the *Escherichia* genus (taxid 561), with *E. coli* (taxid 562) sequences excluded, using the NCBI RefSeq database. Only *Escherichia marmotae* and *Escherichia albertii* were found to possess the *eefRABCD* operon (Fig. S1) while *E. fergusonii, Escherichia ruysiae*, *Escherichia vulneris, and E. hermannii* did not encode it.

Due to the genetic similarity of *Shigella* and *E. coli* (van den Beld and Reubsaet, 2012, Brenner et al., 1972), the presence of the *eef* operon across *Shigella spp*. was investigated. Gene fragments were detected in a small number of *S. boydii* and *S. flexneri* assemblies but only *S. dysenteriae* was found to consistently possess the operon (Fig. S2, Table S5). However, whilst genes belonging to the *eefRABCD* operon were detected in 363 of the 486 *S. dysenteriae* assemblies included in this study, only a single assembly was positive for *eefR.* Moreover, whilst *eefA* was present in all assemblies possessing the operon, sequence coverage averaged at 24.6%. Assembly annotation identified the presence of an insertion sequence in the place of *eefR* and *eefA* (Fig. S3), explaining the absence and truncation of *eefR* and *eefA* respectively, across the *S. dysenteriae* assemblies. In comparison, no insertion sequences were detected within the *eef* operon of *E. coli* ATCC 25922, nor in the 10,000 base pairs up- or down-stream of the operon.

As the EefABC efflux pump has been reported in both *K. aerogenes* and *K. pneumoniae* (Coudeyras et al., 2008, Masi et al., 2005) with differing operon architecture (Fig. S4), the presence of the operon across related Gammaproteobacterial genera was determined. Only low homology orthologs of EefA and EefB were identified in *Yersinia* and *Serratia* while no conserved homologues were identified in *Salmonella*, *Acinetobacter*, *Pseudomonas* or *Photorhabdus* (Table S6). In *Enterobacter,* conservation of the operon differed between species (Fig. S5). Some species, such as *E. chengduensis*, encoded *eefRABC,* but not *eefD*, however sequence identity was only 73% (blastn) indicating that the *E. coli* and *Enterobacter eef* operons are not homologous, and instead the association is likely due to historical literature and gene nomenclature.

### Sequence and structural prediction analysis reveals unique features of the EefABC pump

The *eefABC* operon appears to encode a tripartite RND-efflux pump similar to the AcrAB-TolC assembly in *E. coli*. As there is no experimental structural information currently available on any of the components of EefABC, we conducted sequence analysis to identify potential structural templates and subsequently performed homology modelling using the highest scoring templates.

Comparison with other OMFs of known structure, revealed the closest relatives within the wider OMF-family to be the *Pseudomonas* proteins OprM and OprJ (Fig. S6), followed by CusC, and hence OprM/J were used as structural templates for homology modelling (Fig. S7 and S8 and supplementary text). Structural alignment of EefC with OprM/J produced an alignment with very few gaps allowing for direct mapping of the aligned sequences of EefC onto OprM/J. EefC has significantly shorter extracellular loops in comparison to TolC, in particular the L2 loop which occludes TolC opening (Vaccaro et al., 2008), suggesting a more-open state of the EefC channel (Fig. 4a). Such OMF loops are prominent targets of protective antibodies (Domínguez-Medina et al., 2020) and the non-protruding loops may serve to avoid antibody restriction and LPS occlusion. Additionally, the N-terminal tail of EefC is 30 residues longer than that seen in TolC, making it more similar to the structure of OprN/J (Fig. S8 and Fig. S9).

**Figure 4.**
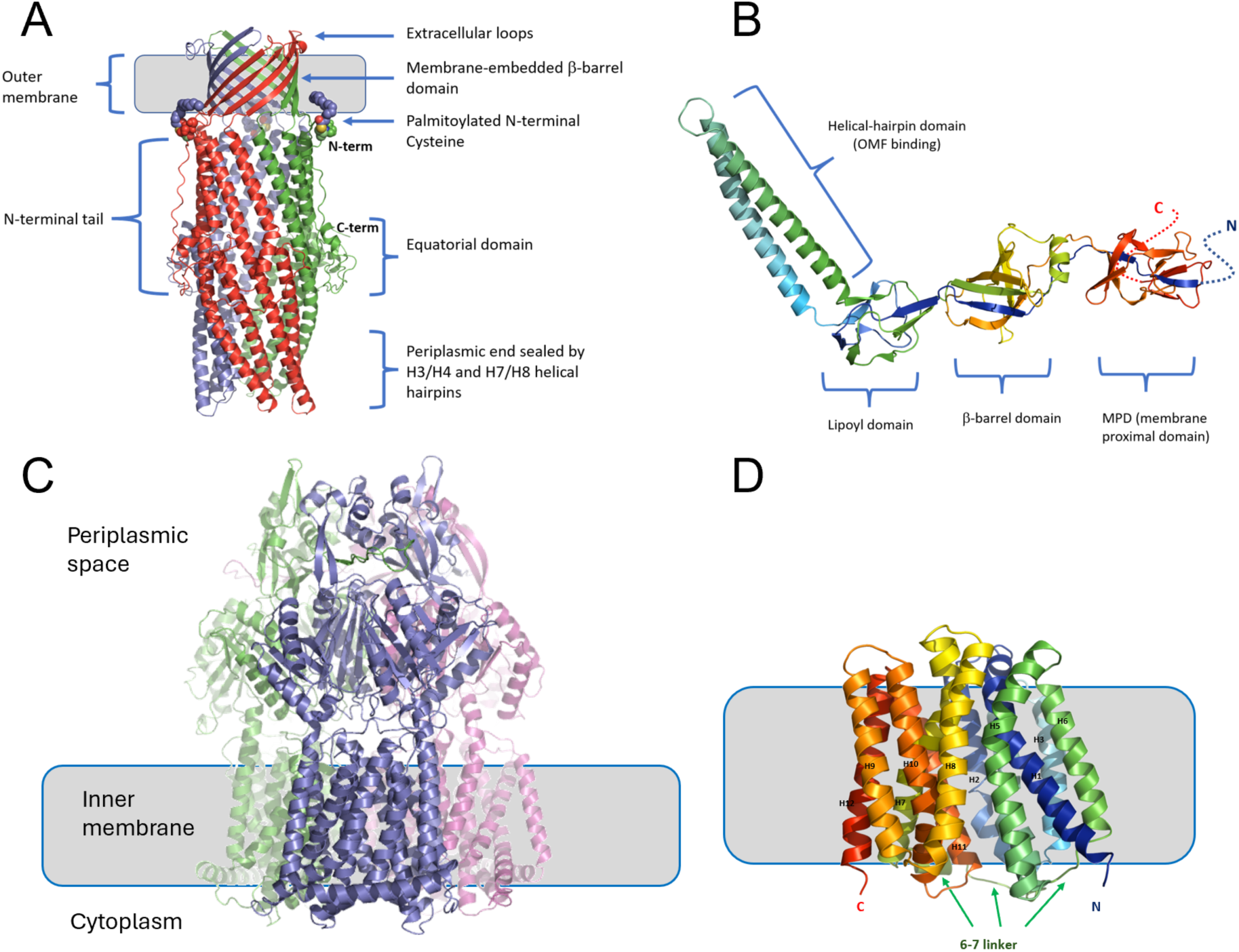
Multi-panel representation of the Eef components. Not to scale.

The comparison of the gating-loop regions in EefC and TolC reveals that they are highly divergent, indicating a locking mechanism markedly different from that observed in TolC, and hence EefC is unlikely to function with any PAPs that normally pair with TolC, and likely only interacts with its cognate PAP EefA. Specifically, several key residues in the helix7/helix8 hairpin of TolC, responsible for gating the TolC channel are different in EefC, e.g. the R367 (TolC), forming part of the so-called “primary gate” that seals the TolC channel by binding to the conserved D153 (TolC; D206 in EefC) and thus anchoring to the helix 4 (Andersen et al., 2002a, Bavro et al., 2008), in EefC is substituted by a small non-polar residue G412, making such interaction impossible (Fig. S8).

In addition, the electrostatic properties of the EefC channel are predicted to be dramatically different to TolC and other OMFs, which is likely to impact heavily the ion and drug selectivity of the channel (Marshall and Bavro, 2020). Firstly, the so-called “secondary gate” of the channel, formed of the prominently conserved double aspartate ring (D371; D373) which forms the basis of cation-selectivity in TolC (Andersen et al., 2002b, Schulz and Kleinekathöfer, 2009), is fully absent in EefC and instead is substituted by small-hydroxylated residues (T416 and T419), a feature which appears to be unique to EefC. Secondly, EefC possesses additional bulky aliphatic residues L415 and L418, which effectively hydrophobically seal the periplasmic end of the channel (Fig. S8).

The PAP EefA was analysed in a similar way, revealing a sequence identity of 53.85% to the *Pseudomonas* MexA; but only 50.13% against the *E. coli* AcrA. Due to the availability of more complete full-length experimental templates, EefA homology models were created using both MexA (2v4d.pdb; (Symmons et al., 2009)) and AcrA (5v5s.pdb; (Wang et al., 2017)) as templates, (utilising both I-TASSER and Swiss-model tools) (Fig. 4b; Fig. S10). While neither of the models delivered the same level of confidence as those for EefC, the alignments with the known PAP structures are readily interpretable, allowing identification of the protein features. A detailed discussion of the EefA structure is given in the Supplementary text with the major findings summarised here.

Alignment of EefA and AcrA results in a direct amino-acid match with only 2 gaps in the alignment, one in the unstructured N-terminal tail, and another at position Q221 (EefA), which has a 4-residue-long deletion relative to AcrA (Fig. 4b; Fig. S10). This region corresponds to the C-terminal end of α-helix 3, which is flanking the β-barrel domain in PAPs, however it is not predicted to affect RND-binding (McNeil et al., 2019). Despite overall similarity with MexA/AcrA there are distinctive differences in the organisation of the EefA, notably in its α-hairpin domain (Fig. 4b; Fig. S11). While the RLS(D) motif, which is thought to be critical for PAP-OMF interaction (Kim et al., 2010, Song et al., 2014, Alav et al., 2021) appears to be preserved in EefA, (R120/L124/S131/D136), there are a number of significant changes in the adjacent residues, notably R123 (K131 in AcrA); V125 (L133); D134 (E142), which would likely result in steric clashes that would preclude direct compatibility with TolC, and such residues are likely to be playing a discriminatory role engaging with EefC (Fig. S11).

The RND-transporter component, EefB, is a large integral membrane protein, predicted to form a functional trimer similar to other transporters in the family (Fig. 4c). Alignment of the EefB with other RND transporters of known function including AcrB (*E. coli*); MtrD (*N. gonorrhoeae*); MexB (*P. aeruginosa*); CusA (*Campylobacter jejuni*) and AdeB (*A. baumanii*) revealed that they are highly similar, with EefB being the most similar to AcrB and MexB (57.0% and 56.8% identity respectively), while CusA and MtrD being most divergent (Fig. S12). The MexB structure was used as a template to generate a high-fidelity homology model of EefB, as there were fewer gaps in the alignment (0.5% vs 0.7% for AcrB) and slightly higher residue overlap with EefB (1038 *vs* 1033 residue overlap respectively).

Overall, the alignment of EefB with MexB (and AcrB) produces very few gaps (the longest is 2 residues long), allowing for unequivocal attribution of secondary structure elements (Fig. S12). As can be seen from the side-by-side comparison of the EefB and AcrB, both present a virtually identical architecture (Fig. S13), with the only notable differences being the shortened loop connecting the TM helices α16 and α17 in EefB (residues 498-507); and the shorter C-terminal tail. Consistent with this, the critical proton-relay residues found in MexB D407, D408, K939 and T976 (Guan and Nakae, 2001) are conserved in EefB (D408, D409, K935 and T972 respectively) and the predicted structures of the access and distal binding pockets of EefB suggest a closer relation to the MexB/AcrB than to MexY type transporters.

The last member of the operon is EefD, a member of the MFS family of efflux pumps that according to the Transporter Classification Database (TCDB) belongs to the 2.A.1.2 group of MFS transporters, related to MDR-function, which also includes Bcr/CflA, EmrD and MdfA (Reddy et al., 2012, Quistgaard et al., 2016) (Fig. S14). This resulted in high-confidence homology models (I-TASSER C-score = 1.82), revealing a classic 6+6 transmembrane helical arrangement closely related to the general topology of MdfA, although with significant differences in the substrate binging cavity, suggesting the substrate range will be notably different between EefD and MdfA (Fig. 4d; Fig. S15) (Adler and Bibi, 2002, Heng et al., 2015, Nagarathinam et al., 2018). These discrepancies make predictions of the possible substrates of EefD problematic, although some overlap with MdfA can be expected (Lewinson et al., 2003), e.g. lipophilic cations such as ethidium.

We used the homology models described above to dock the components into a complete tripartite pump using the cryo-EM structures of AcrAB-TolC (Wang et al., 2017) as a guide. EefABC can indeed be assembled using the same architecture with minimal steric clashes, as can be seen in Fig. 5.

**Figure 5.**
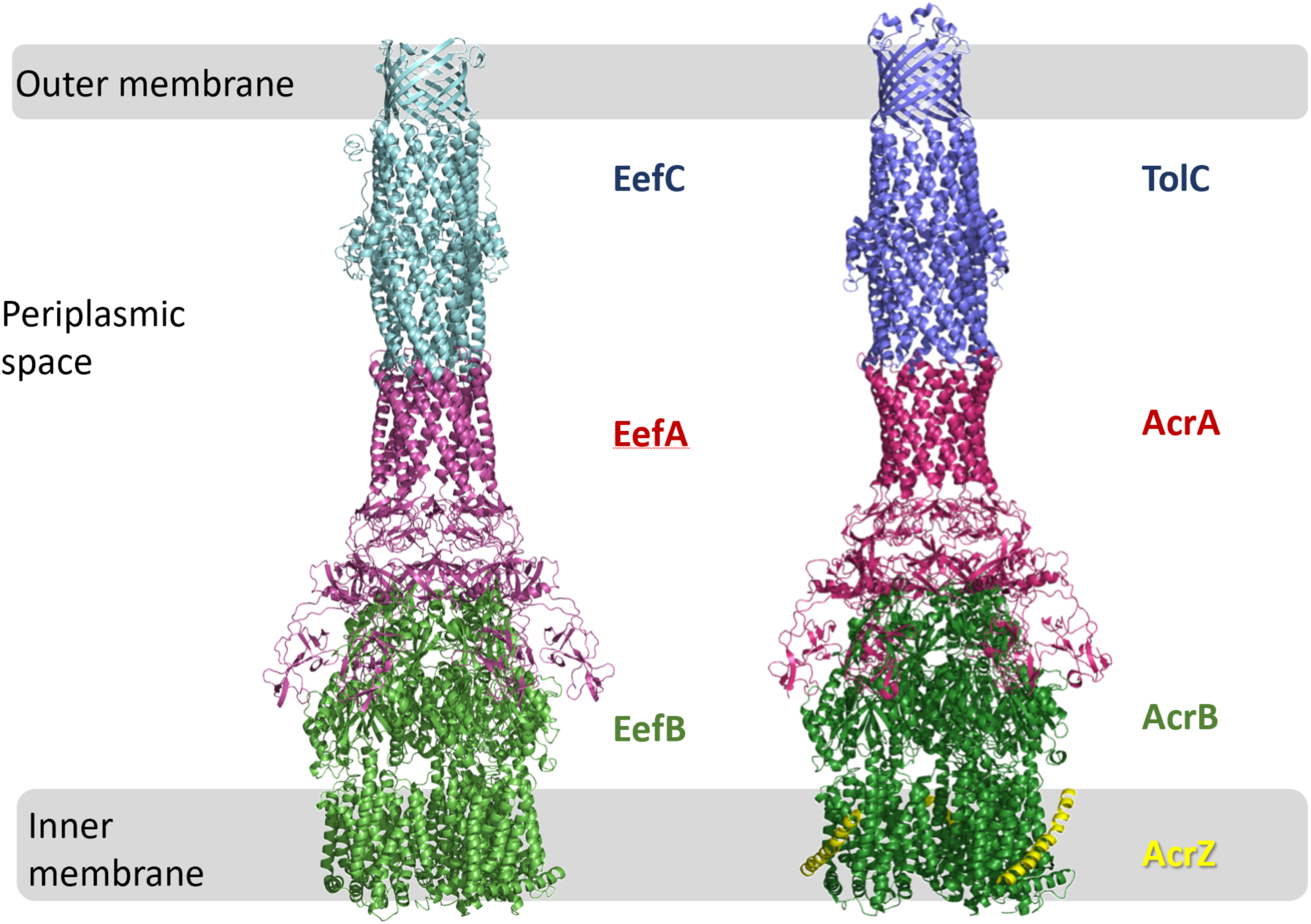
Comparison of the predicted structures of the assembled EefABC and the experimental cryo-EM structure of AcrABZ-TolC. (based on 5066.pdb; (Wang et al., 2017))

### *E. coli* EefABC is not a drug transporter

The high levels of conservation of the *eefRABCD* operon within clinically relevant lineages of *E. coli* suggests it has an important biological function and its substrate profile may differ from other *E. coli* RND pumps due to its unique structure. Antimicrobials are known substrates of RND efflux pumps such as AcrB, and overexpression of RND pumps can confer MDR in both the laboratory and clinic. Therefore, the effect of *eefABC* and *eefD* expression on *E. coli* susceptibility to a range of antimicrobials and dyes was determined. Due to the previously mentioned homology between *E. coli* ATCC 25922 and *E. coli* SMS-3-5, and the well characterised antimicrobial susceptibility profile of ATCC 25922 this strain was used for EefABC characterisation experiments.

Deletion of *eefB* in *E. coli* ATCC 25922 did not increase susceptibility to any antimicrobial or dye tested. Subsequent inactivation of *eefA* and *eefC* to give an *eefABC* knockout also had no effect on the susceptibility of *E. coli* ATCC 25922 to antibiotics (Table 2).

**Table 2.**
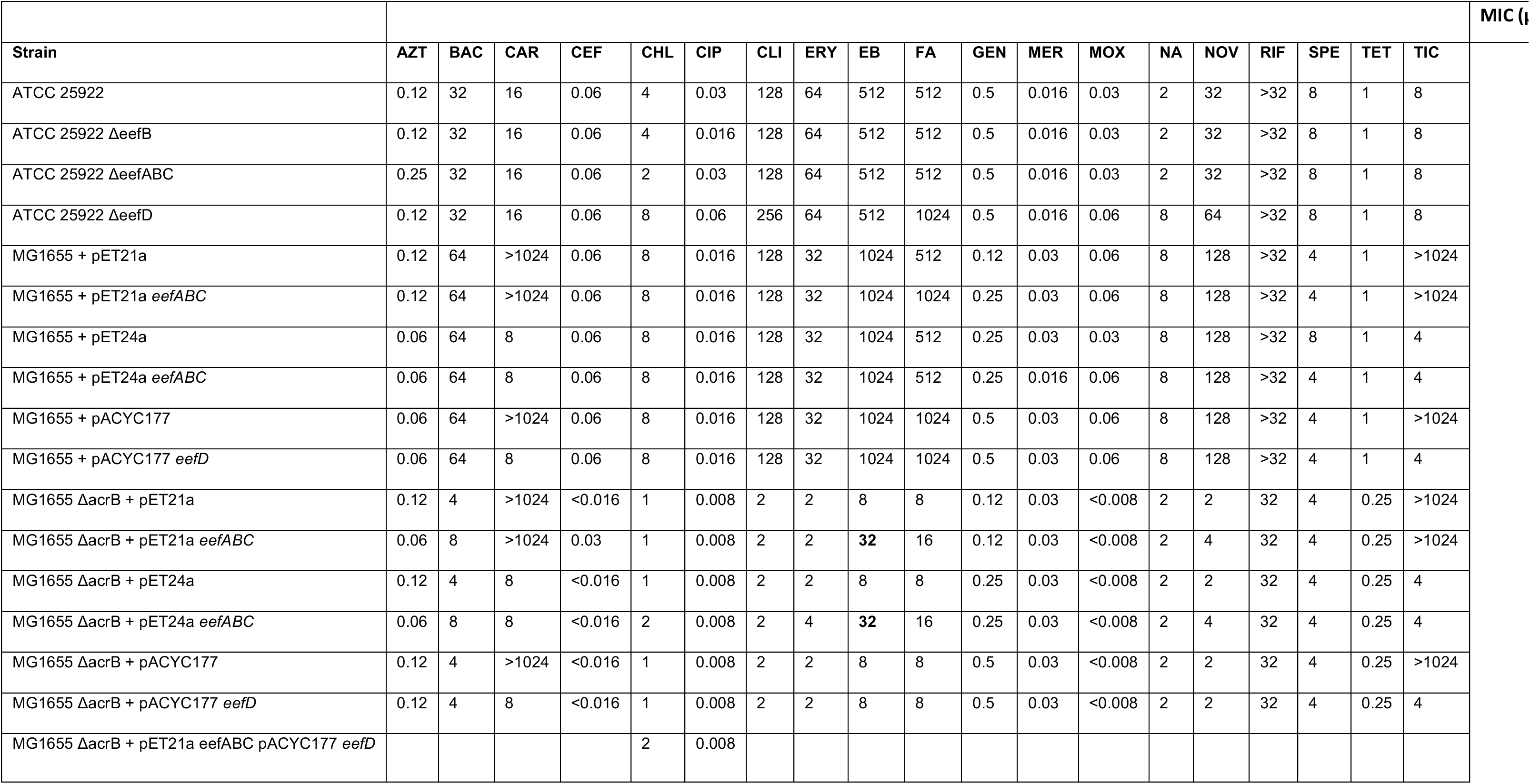
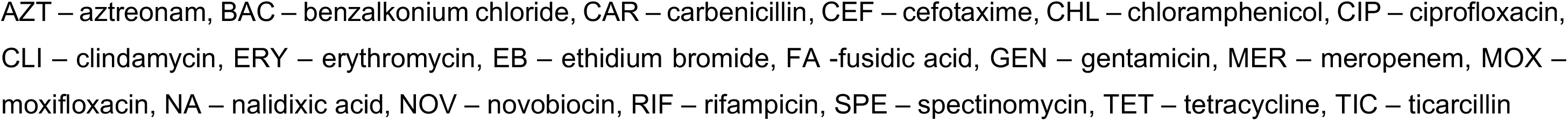
Susceptibility of *E. coli* to antimicrobials following loss and gain of *eefABC* and *eefD*.

|  |  |  |  |  |  |  |  |  |  |  |  |  |  |  |  |  |  |  |  | MIC (μg/ml) |
| --- | --- | --- | --- | --- | --- | --- | --- | --- | --- | --- | --- | --- | --- | --- | --- | --- | --- | --- | --- | --- |
| Strain | AZT | BAC | CAR | CEF | CHL | CIP | CLI | ERY | EB | FA | GEN | MER | MOX | NA | NOV | RIF | SPE | TET | TIC |  |
| ATCC 25922 | 0.12 | 32 | 16 | 0.06 | 4 | 0.03 | 128 | 64 | 512 | 512 | 0.5 | 0.016 | 0.03 | 2 | 32 | >32 | 8 | 1 | 8 |  |
| ATCC 25922 ΔeefB | 0.12 | 32 | 16 | 0.06 | 4 | 0.016 | 128 | 64 | 512 | 512 | 0.5 | 0.016 | 0.03 | 2 | 32 | >32 | 8 | 1 | 8 | |
| ATCC 25922 ΔeefABC | 0.25 | 32 | 16 | 0.06 | 2 | 0.03 | 128 | 64 | 512 | 512 | 0.5 | 0.016 | 0.03 | 2 | 32 | >32 | 8 | 1 | 8 | |
| ATCC 25922 ΔeefD | 0.12 | 32 | 16 | 0.06 | 8 | 0.06 | 256 | 64 | 512 | 1024 | 0.5 | 0.016 | 0.06 | 8 | 64 | >32 | 8 | 1 | 8 | |
| MG1655 + pET21a | 0.12 | 64 | >1024 | 0.06 | 8 | 0.016 | 128 | 32 | 1024 | 512 | 0.12 | 0.03 | 0.06 | 8 | 128 | >32 | 4 | 1 | >1024 |  |
| MG1655 + pET21a *eefABC* | 0.12 | 64 | >1024 | 0.06 | 8 | 0.016 | 128 | 32 | 1024 | 1024 | 0.25 | 0.03 | 0.06 | 8 | 128 | >32 | 4 | 1 | >1024 |  |
| MG1655 + pET24a | 0.06 | 64 | 8 | 0.06 | 8 | 0.016 | 128 | 32 | 1024 | 512 | 0.25 | 0.03 | 0.03 | 8 | 128 | >32 | 8 | 1 | 4 |  |
| MG1655 + pET24a *eefABC* | 0.06 | 64 | 8 | 0.06 | 8 | 0.016 | 128 | 32 | 1024 | 512 | 0.25 | 0.016 | 0.06 | 8 | 128 | >32 | 4 | 1 | 4 |  |
| MG1655 + pACYC177 | 0.06 | 64 | >1024 | 0.06 | 8 | 0.016 | 128 | 32 | 1024 | 1024 | 0.5 | 0.03 | 0.06 | 8 | 128 | >32 | 4 | 1 | >1024 |  |
| MG1655 + pACYC177 *eefD* | 0.06 | 64 | 8 | 0.06 | 8 | 0.016 | 128 | 32 | 1024 | 1024 | 0.5 | 0.03 | 0.06 | 8 | 128 | >32 | 4 | 1 | 4 |  |
| MG1655 ΔacrB + pET21a | 0.12 | 4 | >1024 | <0.016 | 1 | 0.008 | 2 | 2 | 8 | 8 | 0.12 | 0.03 | <0.008 | 2 | 2 | 32 | 4 | 0.25 | >1024 | |
| MG1655 ΔacrB + pET21a *eefABC* | 0.06 | 8 | >1024 | 0.03 | 1 | 0.008 | 2 | 2 | **32** | 16 | 0.12 | 0.03 | <0.008 | 2 | 4 | 32 | 4 | 0.25 | >1024 | |
| MG1655 ΔacrB + pET24a | 0.12 | 4 | 8 | <0.016 | 1 | 0.008 | 2 | 2 | 8 | 8 | 0.25 | 0.03 | <0.008 | 2 | 2 | 32 | 4 | 0.25 | 4 | |
| MG1655 ΔacrB + pET24a *eefABC* | 0.06 | 8 | 8 | <0.016 | 2 | 0.008 | 2 | 4 | **32** | 16 | 0.25 | 0.03 | <0.008 | 2 | 4 | 32 | 4 | 0.25 | 4 | |
| MG1655 ΔacrB + pACYC177 | 0.12 | 4 | >1024 | <0.016 | 1 | 0.008 | 2 | 2 | 8 | 8 | 0.5 | 0.03 | <0.008 | 2 | 2 | 32 | 4 | 0.25 | >1024 | |
| MG1655 ΔacrB + pACYC177 *eefD* | 0.12 | 4 | 8 | <0.016 | 1 | 0.008 | 2 | 2 | 8 | 8 | 0.5 | 0.03 | <0.008 | 2 | 2 | 32 | 4 | 0.25 | 4 | |
| MG1655 ΔacrB + pET21a *eefABC* pACYC177 *eefD* | | | | | 2 | 0.008 | | | | | | | | | | | | | | |

AZT – aztreonam, BAC – benzalkonium chloride, CAR – carbenicillin, CEF – cefotaxime, CHL – chloramphenicol, CIP – ciprofloxacin, CLI – clindamycin, ERY – erythromycin, EB – ethidium bromide, FA -fusidic acid, GEN – gentamicin, MER – meropenem, MOX – moxifloxacin, NA – nalidixic acid, NOV – novobiocin, RIF – rifampicin, SPE – spectinomycin, TET – tetracycline, TIC – ticarcillin

As loss of EefABC function did not alter the drug susceptibility profile of ATCC 25922, *eefABC* was cloned into the pET21a and pET24a plasmids and expressed in *E. coli* MG1655, which does not naturally encode the system. Expression of *eefABC* in *E. coli* MG1655 did not alter susceptibility to any of the antimicrobials tested. AcrB is the dominant RND pump in *E. coli* and this can mask phenotypic changes associated with other efflux systems so it was deleted. However, expression of *eefABC* in the absence of *acrB* still did not reveal any significant changes in susceptibility but did result in a decrease in susceptibility to ethidium bromide and rhodamine 6G suggesting these dyes can be transported by the pump (Table S7).

The MFS pump EefD was also characterised, however both inactivation in *E. coli* ATCC 25922 (loss of function) and expression in *E. coli* MG1655 (gain of function) had no effect on antimicrobial and dye susceptibilities. Expression of *eefD* alone in MG1655 Δ*acrB* was also not found to affect MICs.

A subset of the RND pumps (including CusABC) are known to pump metal ions and form the subfamily of heavy metal efflux (HME)-pumps (Gupta et al., 1999, Long et al., 2012, Lecointre et al., 1998, Klenotic et al., 2020). Due to the structural similarity of EefC to CusC, the susceptibility to heavy metals was also measured. Yet when either *eefB* or *eefABC* were deleted from ATCC 25922, the susceptibility to cobalt, nickel, zinc and iron were not significantly different to the wild-type ATCC 25922.

### EefABC can export ethidium bromide

To confirm the cloned EefABC pump is functional, and that ethidium bromide is a substrate, the intracellular accumulation and efflux rate of ethidium bromide was measured. Expression of EefABC with EefD in the absence of AcrB significantly decreased intracellular accumulation of EtBr and increased the rate at which EtBr was pumped out of cells (Fig. S16 and S7). In addition, deletion of *eefD* significantly slowed efflux of ethidium bromide, which together, shows this inner membrane pump is needed for transport of the substrate across the inner membrane.

## Discussion

The number of RND pumps present in different Gram-negative bacterial species varies and there is increasing evidence that prevalence of RND efflux pumps can also vary between isolates of the same species (Darby et al., 2023, Leus et al., 2020, Nowak et al., 2015, Nemec et al., 2007, Wieczorek et al., 2013). Yet it is still broadly assumed that all *E. coli* isolates possess six RND efflux pumps despite recent work showing that not all six pumps are always functional (Anes et al., 2015, Pugh et al., 2023). Here we further demonstrate that the assumption of six RND pumps is inaccurate; STs belonging to the phylogroups of *E. coli* that are most commonly associated with invasive infection (B2, D, E, F and G) encode a seventh, highly conserved RND pump operon (*eefRABCD*), while the operon was completely absent from phylogroups A, B1 and C, which are generally associated with environmental or commensal lifestyles. The level of conservation suggests that EefRABCD has a biologically important function resulting in a high degree of selection pressure while the distribution in phylogroups commonly associated with infection suggests the system could have a role in infection or survival in the host environment.

Across phylogroups A, B1 and C a 354-nucleotide ORF is found in place of *eefRABCD.* This putative gene *ycjD* runs in the opposite orientation to *eefRABCD* and is highly conserved between STs of these phylogroups. This putative gene is also present in ST69 and ST182 which belong to phylogroup D and E respectively though in these two STs a 12-nucleotide insertion is present at the 3’ end of the gene. Studies from another group support the hypothesis that transcriptional activity is happening at this ORF as public data from their transcriptional start site, term-seq and ribosomal profiling studies show antisense transcriptional and translational activity at the 5’ end of *ycjD* gene (Adams et al., 2021, Thomason et al., 2015, Weaver et al., 2019). It is worth noting, that while *ycjD* is indeed coding a domain of unknown function (DUF), the UniproKB AlphaFold structural prediction for it strongly suggests that it is linked to DNA modification/restriction, as it belongs to the endonuclease/DNA methylase fold.

The EefABC efflux pump was first described in the opportunistic human pathogen *K. aerogenes* and subsequently in *K. pneumoniae*, though operon structure differs between the two *Klebsiella* species (Coudeyras et al., 2008, Masi et al., 2005). In *K. aerogenes* overexpression of *eefABC* decreased susceptibility to erythromycin and ticarcillin (Masi et al., 2006) whilst overexpression of *K. pneumoniae eefA* and *eefB* in an *E. coli* (*ΔacrAB, ΔydhE*) background decreased susceptibility to oxacillin, erythromycin, novobiocin, acriflavine, ethidium bromide and cholate (Ni et al., 2020). However, in this study, neither gain nor loss of *eefABC* function from *E. coli* altered susceptibility to any antibiotics tested, though over-expression did decrease susceptibility to ethidium bromide. Moreover, expression of *eefABC* (in the absence of AcrB) decreased intracellular accumulation of ethidium bromide demonstrating that the pump is functional. No effect was seen upon inactivation of the pump but this is likely because AcrB was still present which would mask any effect. However, deletion of the inner membrane component *eefD* alone, caused significantly slower EtBr efflux even in the presence of AcrB suggesting it has a significant role in transport of substrates across the inner membrane. The apparent difference in substrate profile between *Klebsiella* and *E. coli* could be due to differences in nucleotide sequence or operon structure subsequently altering the regulation of the operon or function of the orthologous protein. In *K. aerogenes* the operon has been demonstrated to be H-NS silenced in laboratory conditions (Masi et al., 2006, Masi et al., 2005), though due to the genomic location of the *eef* operon in *E. coli* and the resulting proximity of the *eefR* promoter to that of *fabI* which encodes an essential enoyl-[acyl-carrier-protein] reductase it is unlikely that the operon is silenced *in vivo* in *E. coli*.

When looking across the *Escherichia* genus, *eefRABCD* was only present in *E. marmotae* and *E. albertii,* though the operon was identified in *S. dysenteriae,* which has high genetic similarity to *E. coli.* Here, *eefR* was generally absent due to the presence of an insertion sequence in place of *eefR* and *eefA.* A further insertion sequence was identified downstream of *eefD.* It has been shown previously that *Shigella* species have higher numbers of insertion sequences when compared to other Gammaproteobacteria such as *E. coli* (Touchon and Rocha, 2007). High numbers of insertion sequences is linked to recent host-specification as the integration of IS elements into the genomes is associated with early stage genome degradation (Moran and Plague, 2004, Hawkey et al., 2020, Yang et al., 2005). The presence of an insertion sequence in place of *eefR* and *eefA* and downstream of *eefD* may therefore suggest the operon is in the process of being lost or degraded in *S. dysenteriae*. Many Gram-negative bacteria including *Salmonella* sp. (taxid 590) did not encode an ortholog of the Eef system but orthologs of EefA and EefB were identified in *Serratia* and *Yersinia* species, suggesting related systems could be dispersed through the Gammaproteobacteria.

Overall, our analysis of EefABC revealed unexpected similarities to the tripartite pumps OprM-MexAB and OprJ-MexCD in *Pseudomonas* (Masuda et al., 2000), rather than to any endogenous *E. coli* paralogues, which may suggest that these pumps have been acquired *via* a lateral gene transfer event, similar to some other plasmid encoded efflux/secretion systems e.g. the EAEC virulence plasmid pAA2 (Imuta et al., 2008).

Given the close similarities of the EefABC system to both MexAB-OprM and MexCD-OprJ and based on previous profiling of these *Pseudomonas* pumps (Masuda et al., 2000), we initially hypothesised that these pumps could have overlapping substrate ranges. However, despite the overall similarity, our tests did not reveal any direct effect of EefABC on transport of antimicrobials, suggesting alternative function. As *E. coli* also possesses HME-RND systems such as CusABC we also investigated the possibility that EefABC may be involved in metal ion tolerance, but we were unable to identify a metal ion substrate for the efflux pump (Table 8). As mentioned above, the EefC lacks the double aspartate “gates” which have been implicated in the coordination of metal ions in the case of TolC (Higgins et al., 2004, Bavro et al., 2008) (Fig. S8). This alone however could not be used to rule out ion-efflux function, as it has previously been noted (Kulathila et al., 2011) that CusC does not (Alcalde-Rico et al., 2016)by itself show any specific features associated with monovalent ion selectivity, with that function rather being associated with the RND transporter, in that case CusA. Indeed, while CusA presents a clear relay of methionine clusters to bind and export Cu(I) and Ag(I) ions (Delmar et al., 2013), there is no identifiable pattern of such residues in the case of EefB. On the other hand, divalent-ion specific pumps such as ZneA, have more restricted binding sites formed from negatively charged residues (Pak et al., 2013), but again, the residues participating in ion coordination are not conserved in EefB.

Aside from conferring resistance to antimicrobials, RND efflux pumps have myriad other physiological roles involving export of both exogenous and endogenous substrates (Henderson et al., 2021). For example, RND efflux pumps in various Gram-negative bacteria have been linked to virulence (Wang-Kan et al., 2017, Nishino et al., 2006). While we have so far been unable to assign clear function to EefRABCD, given the striking distribution and conservation of *eefRABCD* only in phylogroups of *E. coli* associated with disease it is possible that this pump has roles associated with virulence or survival in the host environment and work is continuing to assign biological function.

## Acknowledgements

We thank Karl Hassan for his insightful comments and suggestions in the viva of Hannah Pugh which helped shape sections of this work.

This work was supported by the BBSRC MiBTP DTP grant awarded to HLP (BB/M01116X/1). EMD was funded by the Wellcome Trust (222386/Z/21/Z).

