## Supplementary information for "The RND efflux pump EefABC is highly conserved within lineages of *E. coli* commonly associated with infection" (from another version of this preprint: version 1, 2024-08-01)

**Table S1. Strains used in this study.**

| Species | Genotype | Marker | Reference |
|---|---|---|---|
| *E. coli* | CGSC - DB Strain K-12 MG1655 | | (Blattner *et al.*, 1997) |
| *E. coli* | K-12 MG1655 Δ*acrB* | | (Wang-Kan *et al.*, 2017) |
| *E. coli* | ATCC 25922 | | PHE |
| *E. coli* | ATCC 25922 *eefB*::*aph* | KanR | This study |
| *E. coli* | ATCC 25922 Δ*eefB* | | This study |
| *E. coli* | ATCC 25922 *eefABC*::*aph* | KanR | This study |
| *E. coli* | ATCC 25922 Δ*eefABC* | | This study |
| *E. coli* | ATCC 25922 *eefD*::*aph* | KanR | This study |
| *E. coli* | ATCC 25922 Δ*eefD* | | This study |
| *E. coli* | K-12 MG1655 pACYC177 | AmpR KanR | This study |
| *E. coli* | K-12 MG1655 + pACYC177 *eefD* | KanR | This study |
| *E. coli* | K-12 MG1655 *ΔacrB* pACYC177 | AmpR KanR | This study |
| *E. coli* | K-12 MG1655 Δ*acrB* pACYC177 *eefD* | KanR | This study |
| *E. coli* | K-12 MG1655 + pET21a | AmpR | This study |
| *E. coli* | K-12 MG1655 + pET24a | KanR | This study |
| *E. coli* | K-12 MG1655 Δ*acrB* pET24a | KanR | This study |
| *E. coli* | K-12 MG1655 + pET21a::*eefABC* | AmpR | This study |
| *E. coli* | K-12 MG1655 + pET24a::*eefAB*C | KanR | This study |
| *E. coli* | K-12 MG1655 Δ*acrB* + pET21a::*eefABC* | AmpR | This study |
| *E. coli* | K-12 MG1655 Δ*acrB* + pET24a::*eefABC* | KanR | This study |
| *E. coli* | K-12 MG1655 Δ*acrB* + pET21a::*eefABC* + pACYC177 *eefD* | AmpR KanR | This study |

**Table S2. Primers used in this study.**

| Name | Description | Primer sequence 5'-3' |
|---|---|---|
| *eefABC* cloning forward | Amplifies *eefABC* from the *E. coli* ATCC 25922 chromosome incorporating the *NdeI* restriction site for pET21a/pET24a ligation | TTGACCATATGAAGTATATAGCAACATCTGTAGTG |
| *eefABC* cloning reverse | Amplifies *eefABC* from the *E. coli* ATCC 25922 chromosome incorporating the *XhoI* restriction site for pET21a/pET24a ligation | GTGGTGCTCGAGTTACTTGAGTGATGACATACCGCCG |
| *eefD* cloning forward | Amplifies *eefD* from the *E. coli* ATCC 25922 chromosome incorporating the *ApaLI* restriction site for pACYC177 ligation | CCCCCCGTGCACATGGCTAGAGTCTCTCTTTCATGGGCATTGATTCTTGG |
| *eefD* cloning reverse | Amplifies *eefD* from the *E. coli* ATCC 25922 chromosome incorporating the *PstI* restriction site for pACYC177 ligation | CCCCCCCTGCAGTTACTTTTGATTGTCGATTCTGGTGAGAGCAAC |
| *eefB* deletion forward | Amplifies the *aph* cassette from pKD4 incorporating *eefB* | AGGCTGTTGAAGTTCAGGCTAATGGAGGCAACGCCTGATGTTTgtgtaggctggagctgcttc |

| | | |
|---|---|---|
| *eefB* deletion reverse | overhangs for interruption of *eefB* on the *E. coli* ATCC 25922 chromosome | ACAACACCAGAAAAATAAGAGAACGACGCAACATAACTTAgggaattagccatggtccat |
| *eefAC* deletion forward | Following the inactivation of *eefB* and removal of the *aph* cassette, these primers amplify the *aph* cassette with *eefA* and *eefC* overhangs from the inactivation of the remaining region | TTCAATGAGGAGGGTATTTCTCACCCTGGAGAAAATAATGgtgtaggctggagctgcttc |
| *eefAC* deletion reverse | | AATGCCCATGAAAGAGAGACTCTAGCCATATTATTCCTTAgggaattagccatggtccat |
| *eefD* deletion forward | Amplifies *aph* from the pKD4 plasmid, enabling the inactivation of *eefD,* independent of *eefABC* | CTGGGCGGCGGTATGTCATCACTCAAGTAAGGAATAATgtgtaggctggagctgcttc |
| *eefD* deletion reverse | | GCACCTCATCCTGACAGGCCCCCCTCAGGATGAGGTGTgggaattagccatggtccat |

**Table S3. The number of *E. coli* assemblies downloaded, the number of assemblies removed due to being duplicate assemblies or due to *eef* being split over multiple contigs, and the final number of assemblies used in this study.**

| Phylogroup | ST | Downloaded assemblies | Duplicates removed | Split operons removed | Final assemblies |
|---|---|---|---|---|---|
| A | ST10 | 1758 | 0 | 1 | 1757 |
| | ST119 | 110 | 0 | 0 | 110 |
| | ST167 | 115 | 0 | 0 | 115 |
| | ST540 | 136 | 14 | 0 | 122 |
| | ST617 | 181 | 0 | 0 | 181 |
| B1 | ST17 | 1884 | 0 | 0 | 1884 |
| | ST21 | 2411 | 0 | 0 | 2411 |
| | ST53 | 5 | 0 | 0 | 5 |
| | ST84 | 2 | 0 | 0 | 2 |
| | ST156 | 210 | 10 | 0 | 200 |
| | ST328 | 54 | 0 | 0 | 54 |
| | ST453 | 142 | 6 | 0 | 136 |
| | ST655 | 856 | 238 | 1 | 617 |
| | ST678 | 116 | 9 | 0 | 107 |
| B2 | ST12 | 479 | 41 | 3 | 435 |
| | ST14 | 46 | 0 | 0 | 46 |
| | ST73 | 916 | 0 | 7 | 909 |
| | ST95 | 751 | 0 | 0 | 751 |
| | ST127 | 231 | 0 | 1 | 230 |
| | ST131 | 3186 | 0 | 30 | 3156 |
| | ST144 | 85 | 0 | 0 | 85 |
| C | ST23 | 384 | 19 | 0 | 365 |
| | ST88 | 454 | 33 | 0 | 421 |
| | ST90 | 124 | 8 | 0 | 116 |
| | ST410 | 596 | 23 | 2 | 571 |
| D | ST38 | 815 | 29 | 10 | 776 |
| | ST69 | 696 | 0 | 0 | 696 |
| | ST405 | 525 | 35 | 9 | 481 |
| | ST963 | 55 | 2 | 0 | 53 |
| E | ST11 | 1490 | 132 | 30 | 1328 |
| | ST182 | 76 | 1 | 3 | 72 |
| | ST350 | 34 | 4 | 1 | 29 |
| | ST1266 | 15 | 0 | 2 | 13 |
| F | ST59 | 175 | 18 | 1 | 156 |
| | ST62 | 217 | 122 | 2 | 93 |
| | ST354 | 266 | 22 | 7 | 237 |
| | ST648 | 379 | 0 | 1 | 378 |
| G | ST738 | 38 | 0 | 0 | 38 |

**Table S4. Presence and subsequent gene coverage and nucleotide identity of *eefRABCD* across four *Shigella* species with reference to the *E. coli* ATCC 25922 (CP009072.1) *eefRABCD* operon**

| Species | *n* | Assemblies positive for an *eef* gene | *eefR* | | | *eefA* | | | *eefB* | | | *eefC* | | | *eefD* | | |
|---|---|---|---|---|---|---|---|---|---|---|---|---|---|---|---|---|---|
| | | | *n* | % ID | % coverage | *n* | % ID | % coverage | *n* | % ID | % coverage | *n* | % ID | % coverage | *n* | % ID | % coverage |
| *Shigella boydii* | 469 | 15 | 14 | 99.22 | 100.00 | 14 | 97.95 | 100.00 | 15 | 96.63 | 62.36 | 9 | 97.32 | 100.00 | 14 | 97.89 | 83.10 |
| *Shigella dysenteriae* | 486 | 363 | 1 | 98.94 | 99.82 | 363 | 97.62 | 26.40 | 363 | 98.29 | 100.00 | 363 | 97.38 | 100.00 | 363 | 97.25 | 98.97 |
| *Shigella flexneri* | 352 | 1 | 1 | 99.12 | 100.00 | 1 | 97.95 | 100.00 | 1 | 98.17 | 100.00 | 1 | 97.53 | 100.00 | 1 | 98.37 | 100.00 |
| *Shigella sonnei* | 360 | 0 | 0 | 0 | 0 | 0 | 0 | 0 | 0 | 0 | 0 | 0 | 0 | 0 | 0 | 0 | 0 |

Percentages shown are means.

### Table S5. Conservation of *eefB* homologs across five closely related bacterial genera.

| | | Genome | | | | | | | | | |
|---|---|---|---|---|---|---|---|---|---|---|---|
| | | 1 | | 2 | | 3 | | 4 | | 5 | |
| Genus | Taxid | Coverage (%) | Identity (%) | Coverage (%) | Identity (%) | Coverage (%) | Identity (%) | Coverage (%) | Identity (%) | Coverage (%) | Identity (%) |
| *Yersinia* | 629 | 98 | 68.47 | 98 | 68.47 | 98 | 68.52 | 98 | 68.52 | 98 | 68.49 |
| *Serratia* | 613 | 99 | 69.76 | 92 | 72.89 | 85 | 72.89 | 92 | 72.82 | 86 | 72.82 |
| *Pseudomonas* | 286 | 13 | 98.08 | 31 | 74.03 | 29 | 75.84 | 30 | 75.63 | 38 | 75.63 |
| *Photorhabdus* | 29487 | 25 | 72.43 | 25 | 72.04 | 25 | 72.04 | 25 | 72.04 | 36 | 71.84 |
| *Salmonella* | 590 | 44 | 74.89 | 47 | 74.89 | 53 | 74.89 | 32 | 76.01 | 36 | 75 |
| *Acinetobacter* | 469 | 98 | 68.49 | 22 | 70.68 | 10 | 77.36 | 10 | 76.76 | 32 | 70.88 |

**Table S6. Presence and subsequent gene coverage of *eefRABCD* across the *E. coli* assemblies with the *eefRABCD* operon of *E. coli* ATCC used as a reference.**

| Phylogroup | ST group | Assemblies (*n*) | Mean sequence coverage to ATCC 25922 *eef* operon (%) | | | | |
|---|---|---|---|---|---|---|---|
| | | | *eefR* | *eefA* | *eefB* | *eefC* | *eefD* |
| A | 10 | 1757 | 0.06 | 0.06 | 0.06 | 0.06 | 0.00 |
| | 119 | 110 | 0.00 | 0.00 | 0.00 | 0.00 | 0.00 |
| | 167 | 115 | 0.00 | 0.00 | 0.00 | 0.00 | 0.00 |
| | 540 | 122 | 0.00 | 0.00 | 0.00 | 0.00 | 0.00 |
| | 617 | 181 | 0.55 | 0.55 | 0.55 | 0.55 | 0.55 |
| B1 | 17 | 1884 | 0.00 | 0.00 | 0.00 | 0.00 | 0.00 |
| | 21 | 2411 | 0.00 | 0.00 | 0.00 | 0.00 | 0.00 |
| | 53 | 5 | 0.00 | 0.00 | 0.00 | 0.00 | 0.00 |
| | 84 | 2 | 0.00 | 0.00 | 0.00 | 0.00 | 0.00 |
| | 156 | 200 | 0.50 | 0.00 | 0.00 | 0.00 | 0.00 |
| | 328 | 54 | 0.00 | 0.00 | 0.00 | 0.00 | 0.00 |
| | 453 | 136 | 0.00 | 0.00 | 0.00 | 0.00 | 0.00 |
| | 655 | 617 | 0.00 | 0.00 | 0.00 | 0.00 | 0.00 |
| | 678 | 107 | 0.00 | 0.00 | 0.00 | 0.00 | 0.00 |
| B2 | 12 | 435 | 100.00 | 99.84 | 99.77 | 99.77 | 100.00 |
| | 14 | 46 | 100.00 | 100.00 | 100.00 | 100.00 | 100.00 |
| | 73 | 909 | 100.00 | 100.00 | 100.00 | 100.00 | 99.97 |
| | 95 | 751 | 99.99 | 100.00 | 100.00 | 100.00 | 100.00 |
| | 127 | 230 | 99.98 | 100.00 | 100.00 | 100.00 | 100.00 |
| | 131 | 3156 | 99.87 | 99.87 | 99.91 | 99.91 | 99.95 |
| | 144 | 85 | 99.99 | 100.00 | 100.00 | 100.00 | 100.00 |
| C | 23 | 365 | 0.27 | 0.27 | 0.08 | 0.00 | 0.00 |
| | 88 | 421 | 0.00 | 0.00 | 0.02 | 0.00 | 0.00 |
| | 90 | 116 | 0.00 | 0.00 | 0.00 | 0.00 | 0.00 |
| | 410 | 571 | 0.00 | 0.00 | 0.00 | 0.00 | 0.00 |
| D | 38 | 776 | 99.69 | 99.74 | 99.74 | 99.74 | 99.74 |
| | 69 | 696 | 0.00 | 0.00 | 0.00 | 0.00 | 0.00 |
| | 405 | 481 | 99.97 | 100.00 | 100.00 | 100.00 | 100.00 |
| | 963 | 53 | 99.99 | 100.00 | 100.00 | 100.00 | 100.00 |
| E | 11 | 1328 | 99.99 | 100.00 | 99.90 | 100.00 | 100.00 |
| | 182 | 72 | 0.00 | 0.00 | 0.00 | 0.00 | 0.00 |
| | 350 | 29 | 99.68 | 100.00 | 100.00 | 100.00 | 100.00 |
| | 1266 | 13 | 100.00 | 100.00 | 100.00 | 100.00 | 100.00 |
| F | 59 | 156 | 99.32 | 99.36 | 99.36 | 99.21 | 99.36 |
| | 62 | 93 | 98.92 | 98.91 | 98.92 | 98.92 | 98.92 |
| | 354 | 237 | 98.50 | 99.07 | 99.16 | 99.16 | 99.35 |
| | 648 | 378 | 100.00 | 100.00 | 100.00 | 100.00 | 100.00 |
| G | 738 | 38 | 100.00 | 100.00 | 100.00 | 100.00 | 100.00 |

**Table S7. Susceptibility of *E. coli* to metals, dyes and bile salts**

| Strain | MIC (µg/mL) | | | | | | | | | | |
|---|---|---|---|---|---|---|---|---|---|---|---|
| | Metals | | | Dyes | | | | Bile Salts | | | ABX |
| | CoCl2 | CuSO4 | NiCl | CV | EB | MB | R6G | SC | SDC | CDC | POL |
| ATCC 25922 | 512 | 256 | 512 | 32 | 512 | >1024 | 1024 | >32,096 | 32,096 | 32,096 | 0.5 |
| ATCC 25922 Δ*eefB* | | | | 32 | 512 | >1024 | 512 | | | | |
| ATCC 25922 Δ*eefABC* | 512 | 256 | 512 | 32 | 512 | >1024 | 512 | >32,096 | 32,096 | 32,096 | 0.5 |
| ATCC 25922 Δ*eefD* | 512 | | | 32 | 512 | >1024 | 1024 | | | | 0.5 |
| MG1655 + pET21a | | | | 16 | 1024 | >1024 | 512 | | | | |
| MG1655 + pET21a *eefABC* | | | | 32 | 1024 | >1024 | 512 | | | | |
| MG1655 + pET24a | | | | 16 | 1024 | >1024 | 512 | | | | |
| MG1655 + pET24a *eefABC* | | | | 16 | 1024 | >1024 | 512 | | | | |
| MG1655 + pACYC177 | | | | 32 | 1024 | >1024 | 512 | | | | |
| MG1655 + pACYC177 *eefD* | | | | 32 | 1024 | >1024 | 512 | | | | |
| MG1655 Δ*acrB* + pET21a | 512 | 256 | 512 | 4 | 8 | 8 | 8 | >32,096 | 16,048 | 8,024 | 1 |
| MG1655 Δ*acrB* + pET21a *eefABC* | 512 | 256 | 512 | 4 | **32** | 16 | **32** | >32,096 | 32,096 | 4,012 | 0.5 |
| MG1655 Δ*acrB* + pET24a | | | | 4 | 8 | 8 | 8 | | | | |
| MG1655 Δ*acrB* + pET24a *eefABC* | | | | 4 | **32** | 16 | **32** | | | | |
| MG1655 Δ*acrB* + pACYC177 | 512 | 526 | 512 | 4 | 8 | 8 | 8 | >32,096 | 16,048 | 4,012 | 0.5 |

| | | | | | | | | | | | |
|---|---|---|---|---|---|---|---|---|---|---|---|
| MG1655 Δ*acrB* + pACYC177 *eefD* | 512 | 256 | 512 | 4 | 8 | 8 | 8 | >32,096 | 16,048 | 4,012 | 0.5 |
| MG1655 Δ*acrB* + pET21a eefABC pACYC177 *eefD* | 512 | 526 | 512 | | | | | >32,096 | 16,048 | 4,012 | 0.5 |

CoCl2 – cobalt chloride CuSO4 – copper sulphate NiCl – nickel chloride CV – crystal violet EB – ethidium bromide MB – methylene blue R6G – rhodamine 6G SC – SDC – CDC – POL -

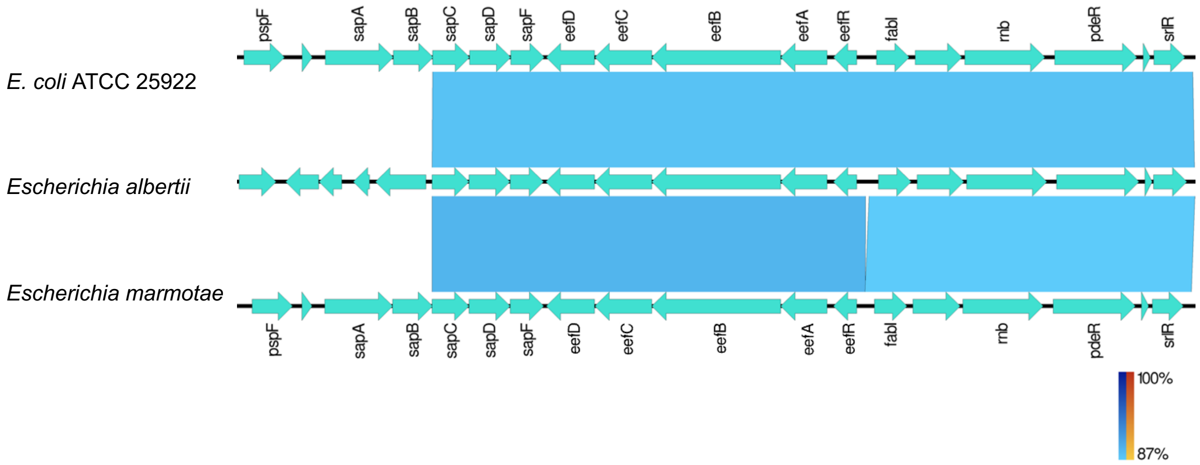


**Figure S1. Alignment of the *eef* operon of multiple *Escherichia* species.**

As *eefRABC* was found to be highly prevalent across *E. coli,* the wider *Escherichia* genus was searched. Only *E. albertii* and *E. marmotae* were found to possess any of the *eef* genes though conservation was significantly lower than when comparing *E. coli* assemblies. Here, *E. albertii* (CP070290.2) and *E. marmotae* (CP056165.1) are aligned to the *eefRABCD* region of *E. coli* ATCC 25922 (CP009072.1) using blastn and visualised using EasyFig. *E. albertii* and *E. marmotae* were annotated using Prokka.

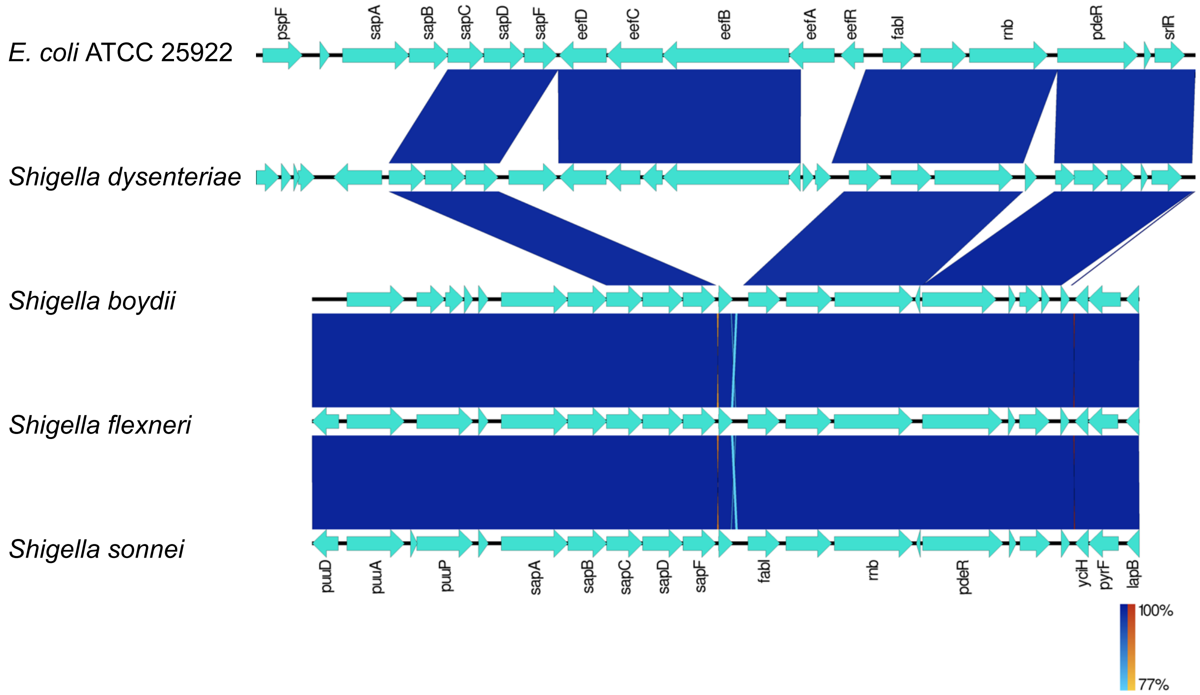


**Figure S2. Alignment of *E. coli* ATCC 25922 *eef* operon and corresponding genome region in multiple *Shigella* species.**

*Shigella* species have high genetic similarity to *E. coli,* therefore we looked for the presence of *eefRABCD* across assemblies of *S. boydii* (*n* = 469), *S. dysenteriae* (*n* = 486), *S. flexneri* (*n* = 352) and *S. sonnei* (*n* = 360). The operon was completely absent from *S. sonnei.* Only one of the *S. flexneri* assemblies possessed at least one *eef* gene, whilst 15 *S. boydii* assemblies had a gene homologous to the *eefRABCD* operon. However, 363 of the 486 *S. dysenteriae* assemblies were positive for genes homologous to at least of the *eefRABCD* genes however, *eefR* was absent from all assemblies except one. Alignment of four NCBI *Shigella* nucleotide sequences with EasyFig demonstrates the high nucleotide homology between *E. coli* ATCC 25922 (CP009072.1) and *S. dysenteriae* (CP026774.1). *S. boydii* (CP026836.1), *S. flexneri* (AE005674.2) and *S. sonnei* (CP055292.1) all lack the operon but are highly homologous to each other.

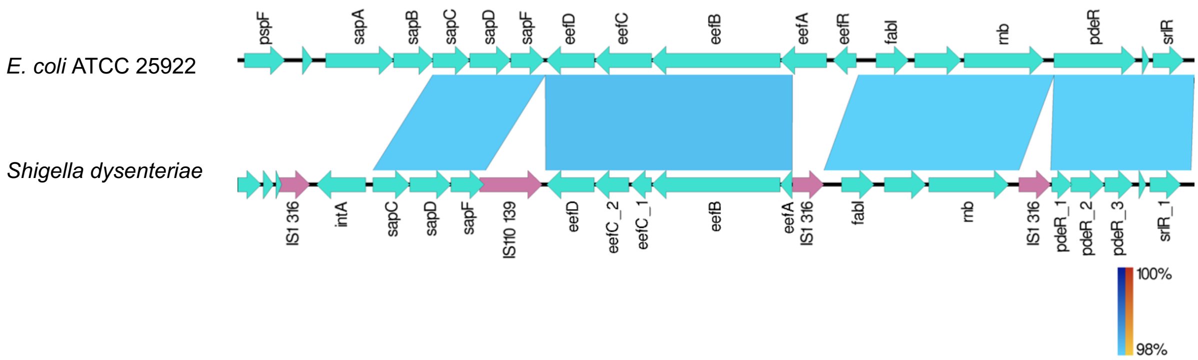


**Figure S3. *S. dysenteriae* possesses insertion sequences within the *eef* operon.** The presence of *eefRABCD* across *S. boydii, S. flexneri, S. dysenteriae* and *S. sonnei* was identified using ABRicate and a custom blast database generated from the *eefRABC* gene sequences of *E. coli* ATCC 25922 (CP009072.1). This data suggested *eefBCD* were present across *S. dysenteriae* but *eefR* and *eefA* were either completely absent or partially absent. Annotation of *S. dysenteriae* (CP026774.1) with ISEScan identified an IS3 insertion sequence in place of *eefR* and *eefA* and a further IS110 element downstream of *eefD. S. dysenteriae* was annotated using Prokka.

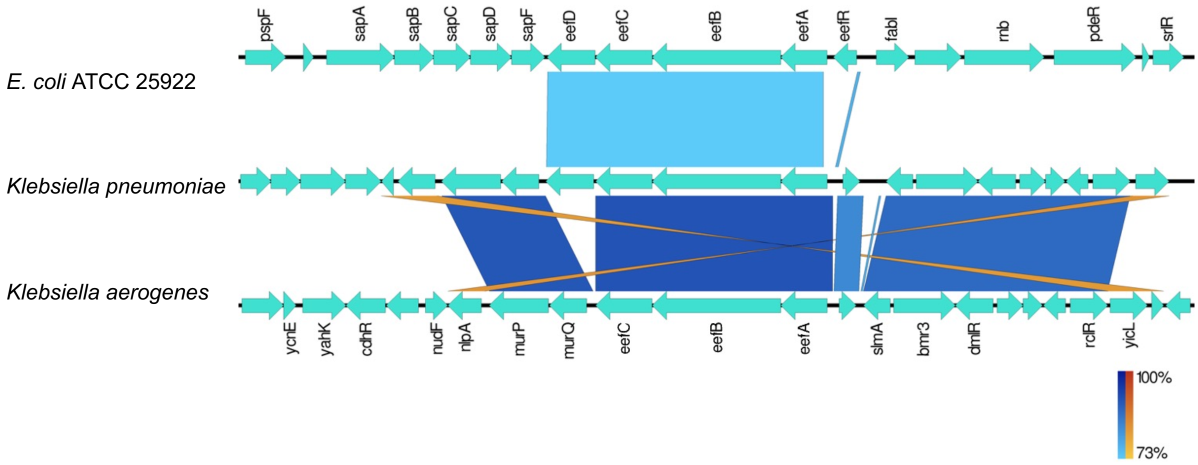


**Figure S4. Alignment of the *E. coli* and *Klebsiella* species *eef* operons**

The presence of *eef* has been reported in both *K. aerogenes* and *K. pnuemoniae* however in *K. aerogenes* the operon was reported to be H-NS silenced (Masi et al., 2005). Alignment of *K. pneumoniae* (NC_016845.1) and *K. aerogenes* (NZ_CP041925.1) against *E. coli* ATCC 25922 (CP009072.1) demonstrates not only differing operon architecture between the *Klebsiella* species but also a different genome location and reduced nucleotide homology to the *E. coli eef* operon. *K. aerogenes* was annotated using Prokka.

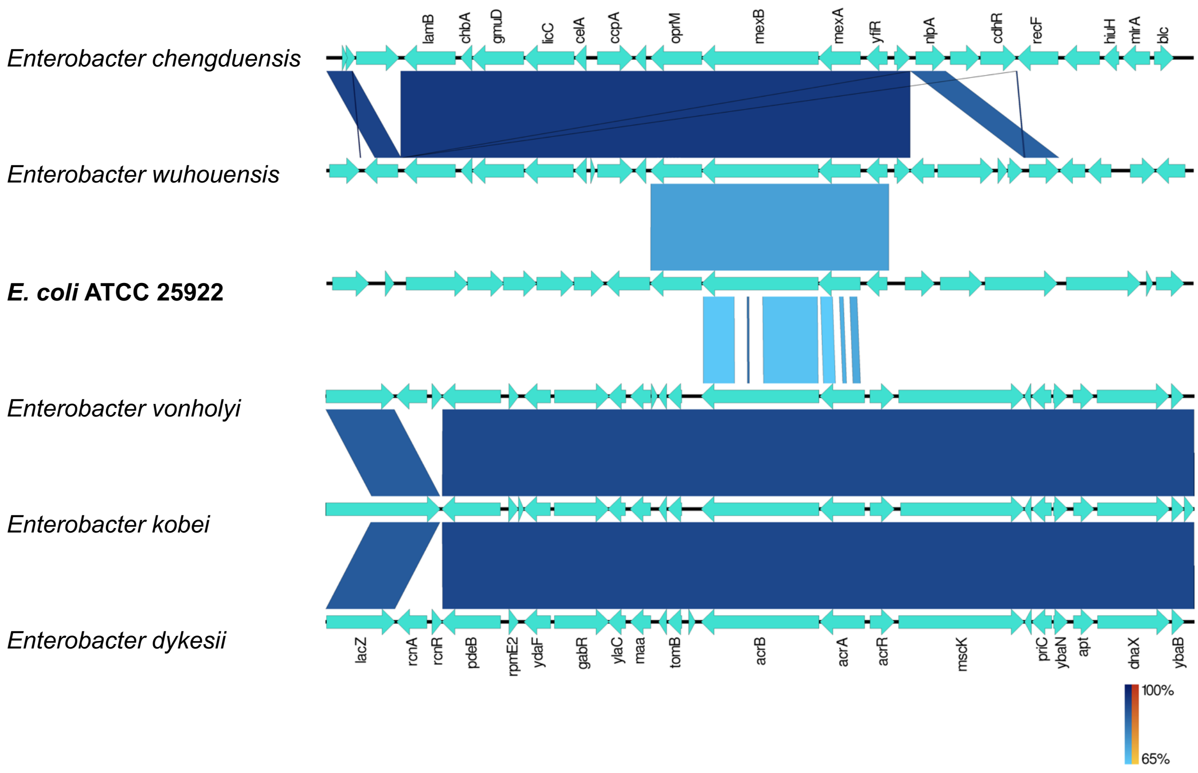


**Figure S5. Alignment of *Enterobacter* species against *E. coli* ATCC 25922.** Though *E. aerogenes* has since been reclassified as a *K. aerogenes* we looked at the *eef* presence in five *Enterobacter* species. Nucleotide blast was used to identify the position of genes with homology in *E. vonholyi* (VTUC01000001.1), *E. dykesii* (VTTY01000003.1), *E. wuhouensis* (SJOO01000006.1), *E. kobei* (KI973153.1) and *E. chengduensis* (CP043318.1) and these genome regions were then aligned and visualised using EasyFig. A four gene operon was identified in *E. chengduensis* but nucleotide identity was low when compared to *E. coli. Enterobacter* sequences were annotated using Prokka.

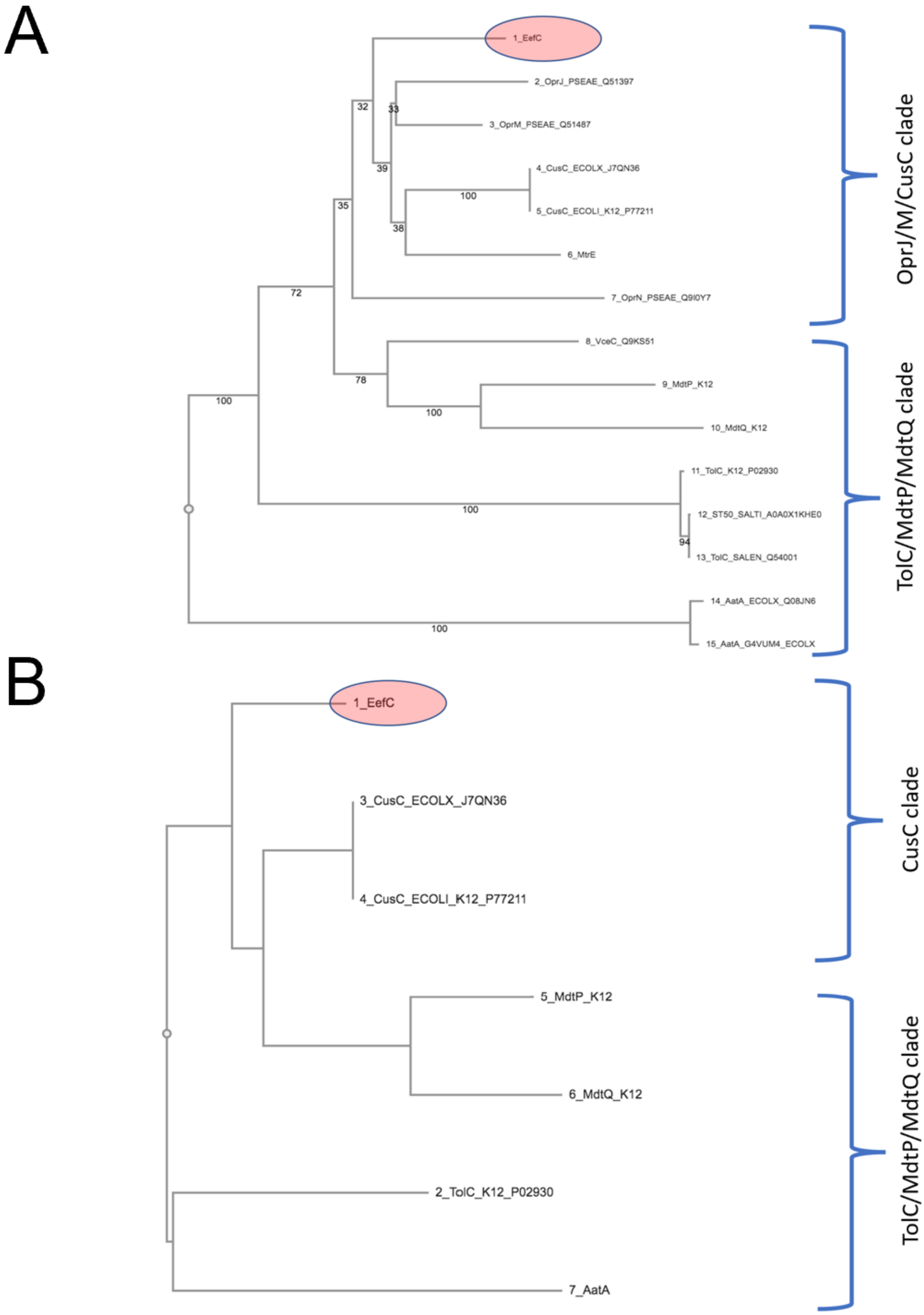


**Figure S6. Dendrograms, showing the phylogenetic connections of EefC to the wider OMF family (A) and its relations with the OMFs of *E. coli* (B).**
Note the clear connection to the OprM/OprJ clade and the CusC branch, justifying their choice as templates for homology modelling.

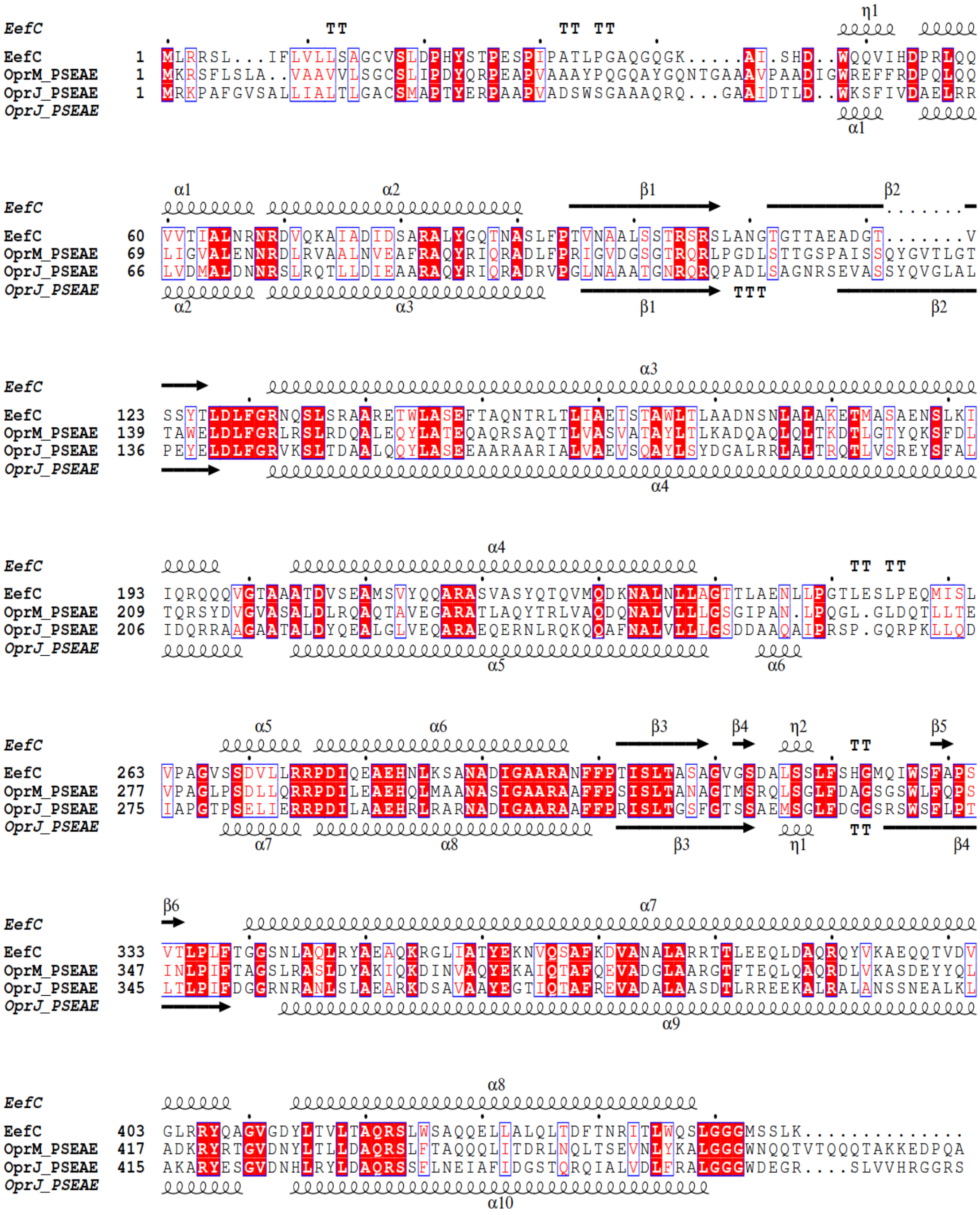

**Figure S7. An MSA of EefC with closely related members of the OMF family**. As can be seen, EefC is closely related to the OprJ/M/N branch of the OMF family, allowing for reliable homology modelling using them as templates

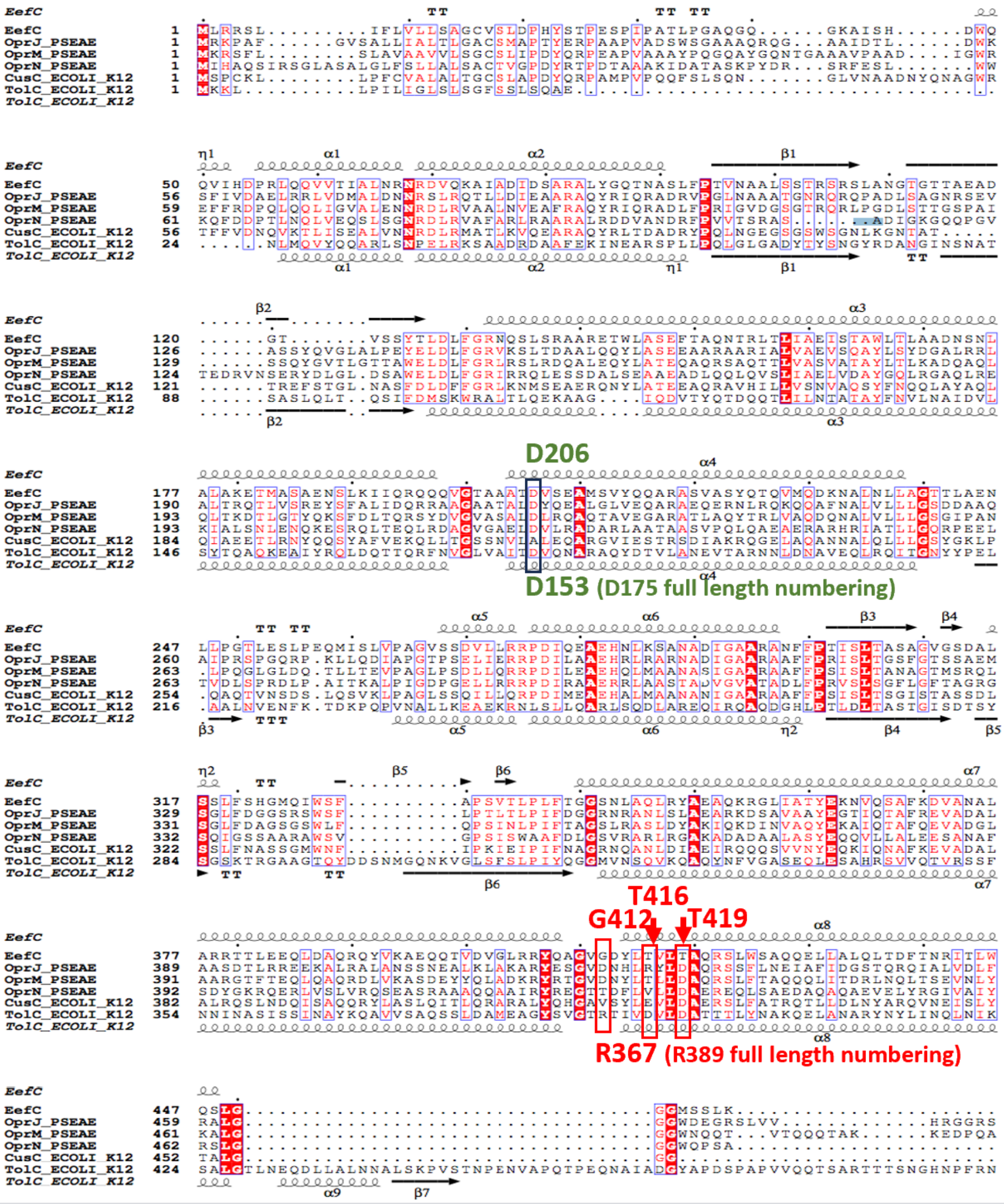


**Figure S8. A wider MSA of OMFs, including TolC, showing the closer association of EefC to the OprJ/M/N clade of OMFs than to TolC, and highlighting some critical differences in gating residues controlling the opening of the EefC vs TolC channels.** Residue numbers are given for full-length, non-processed EefC, while for historical reasons, both full-length and mature protein residues numbers (as per (Koronakis et al., 2000)) are provided for TolC.

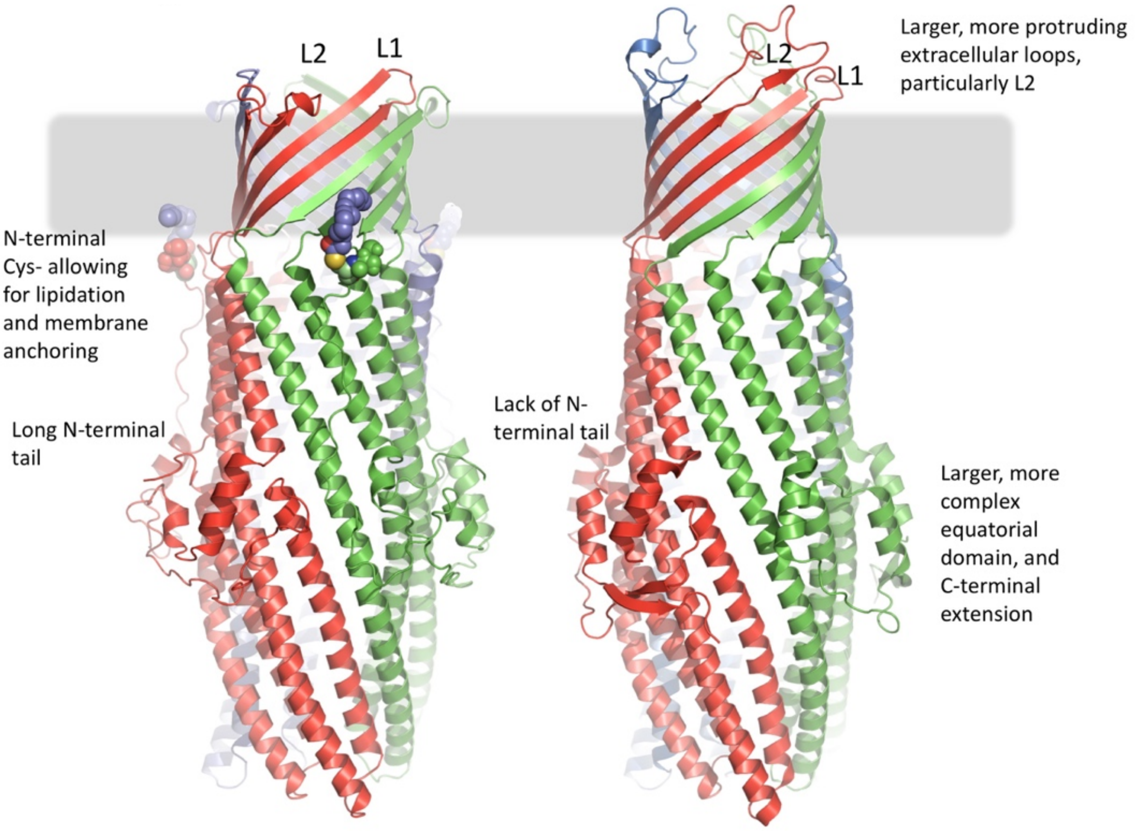


**Figure S9. Key structural differences between EefC (left) and TolC channels as discussed in the main text.**

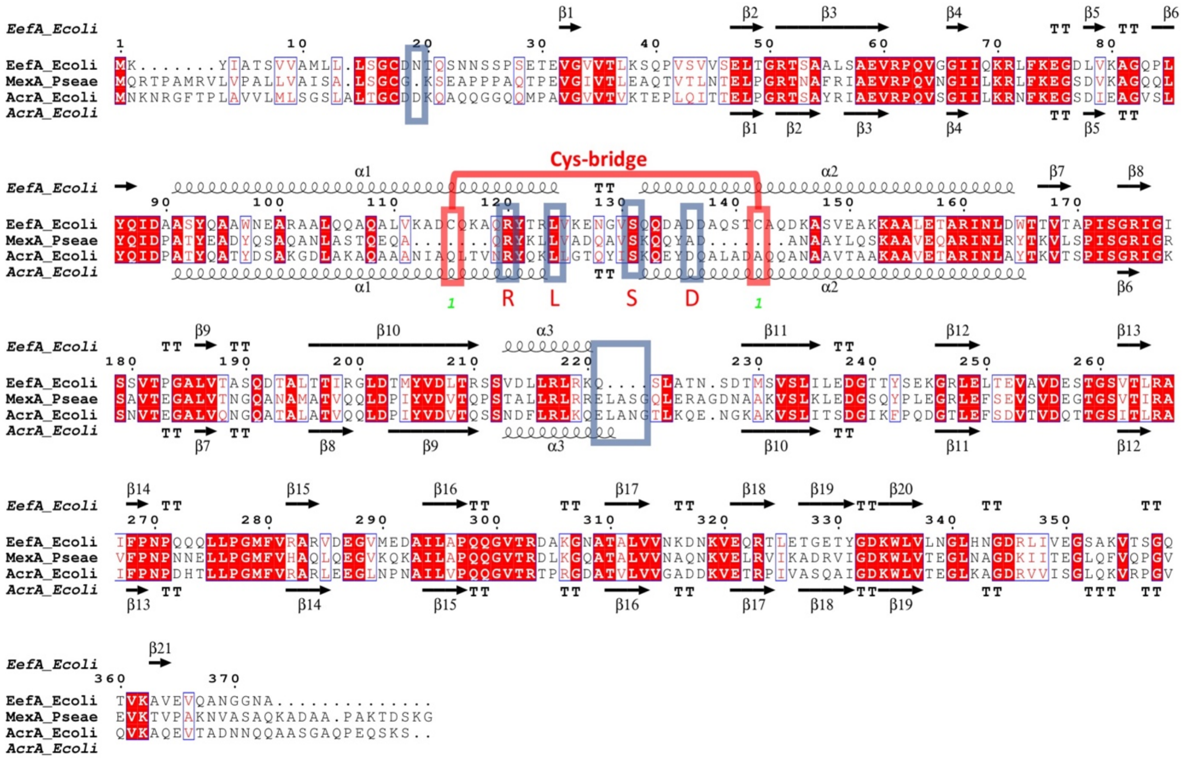


**Figure S10. An MSA of EefA with periplasmic adapter proteins of known structure, used as templates for its structural modelling.** Secondary structure elements are indicated above the and below the alignment for EefA (top) and AcrA, respectively. The RLS motif at the tip of the alpha-helical hairpin domain of the PAPs is highlighted with a set of boxes and letters, as is the 4-residue deletion in EefA relative to the other PAPs as discussed in the text.

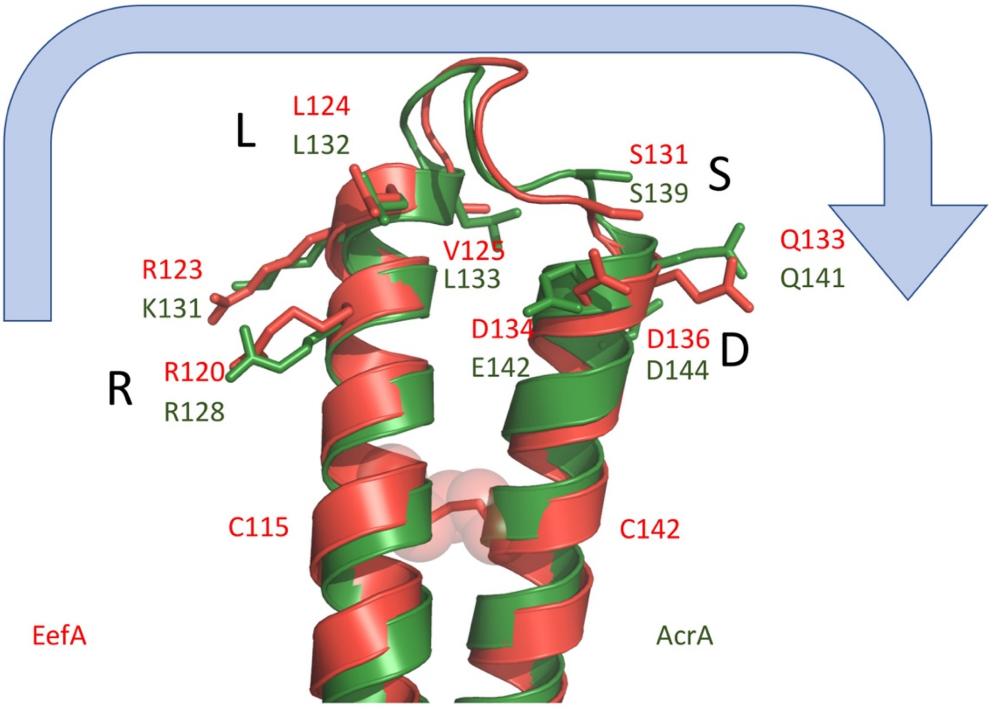


**Figure S11. Residue conservation at the tip of the alpha-helical hairpin domain of EefA vs AcrA suggest radically different OMF-docking patterns, and likely cross-incompatibility between the PAPs and their respective OMFs.**

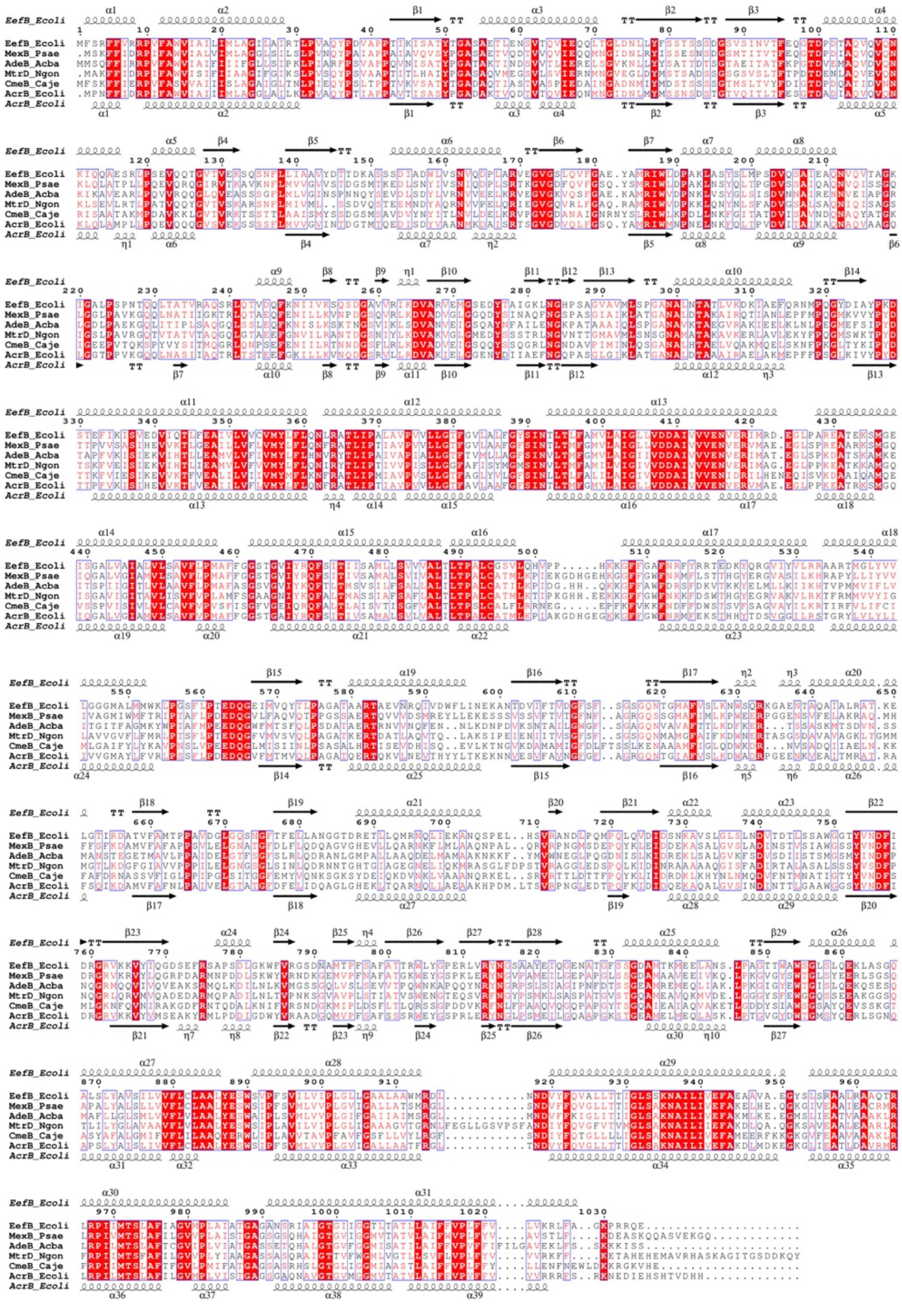
EefB_Ecoli
MexB_Psae
AdeB_Acba
MtrD_Ngon
CmeB_Caje
AcrB_Ecoli

**Figure S12. Structural alignment comparing the predicted organisation of EefB (top) with the empirical structure of *E. coli* AcrB (bottom) and sequences of other RND transporters of known structure.**

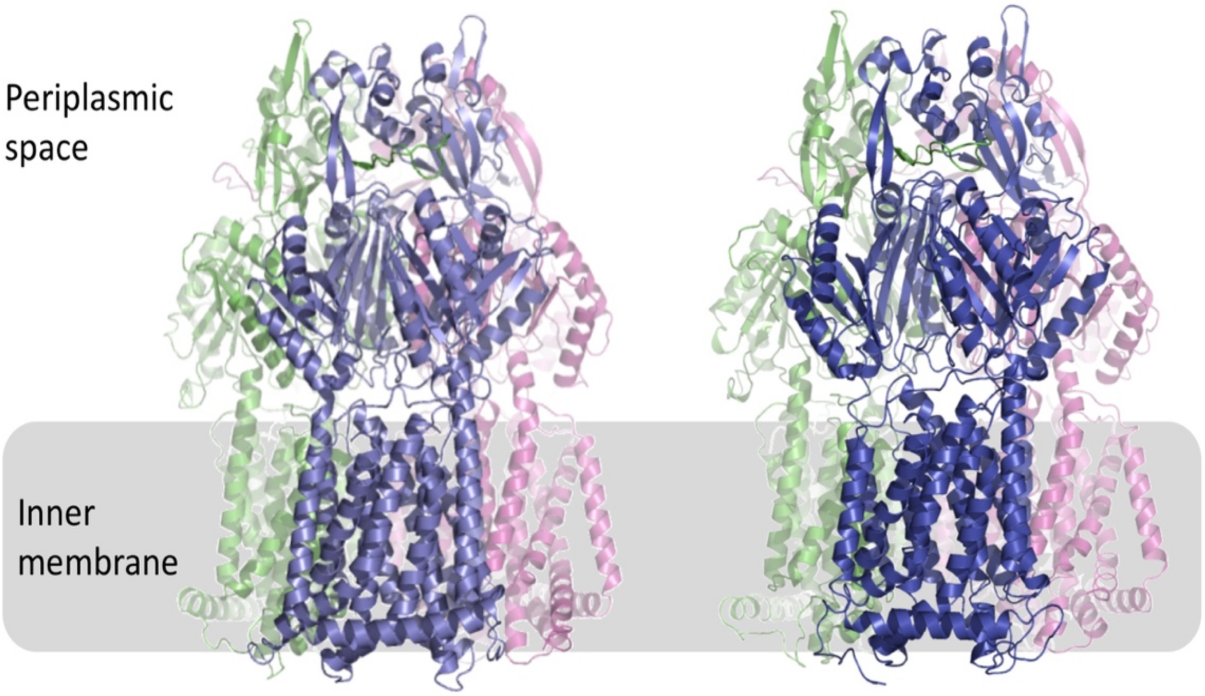


**Figure S13. Comparison of the predicted structures of the assembled EefB trimer (left) and the experimental structure of AcrB (based on 2GIF.pdb) (Seeger et al., 2006).**

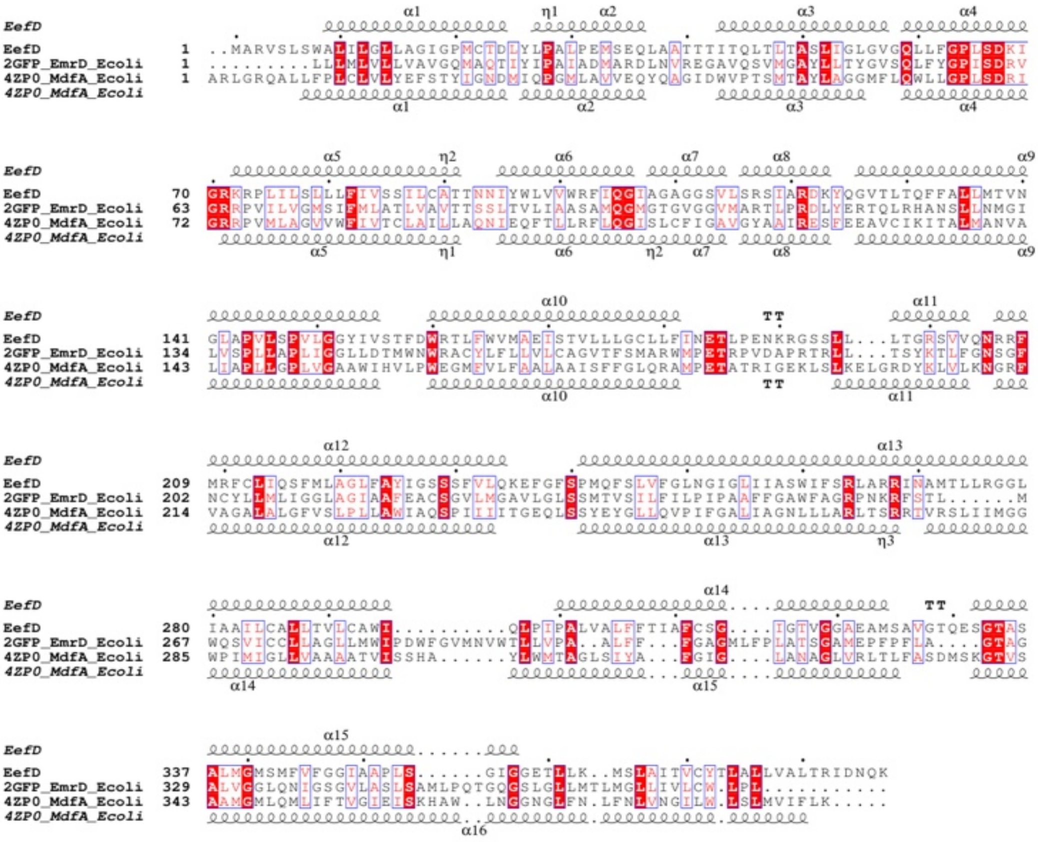

**Figure S14. A MSA comparing the predicted structure of EefD with the empirically determined structures of *E. coli* EmrD and MdfA. Secondary structure elements are shown above and below the alignment.**

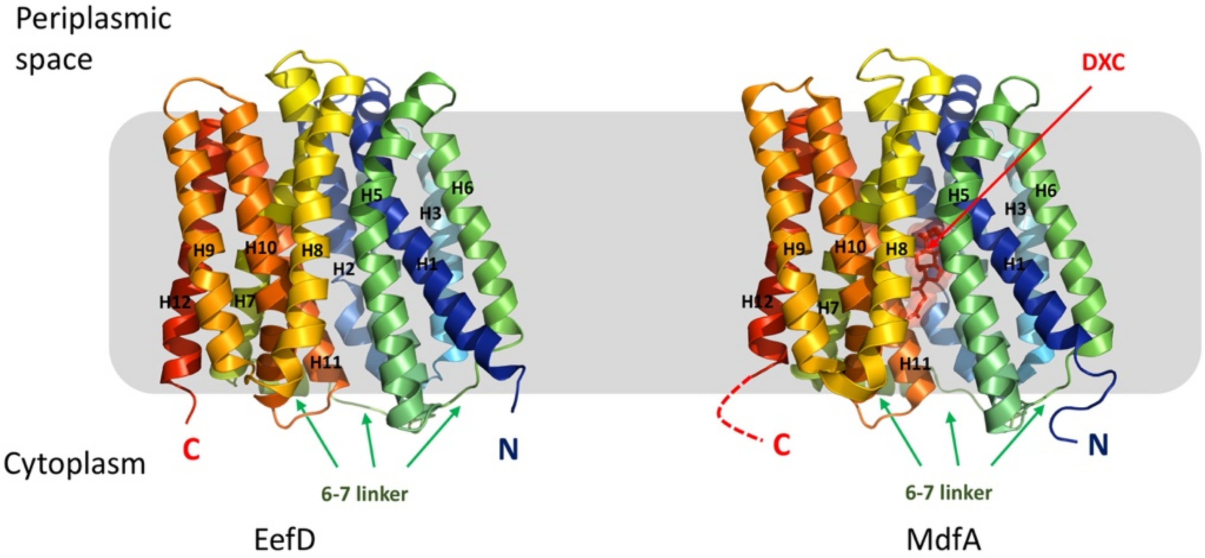


**Figure S15. Side-by-side comparison of the predicted structure of EefD (left, modelled in an inward facing state) with the empirically determined structure of *E. coli* MdfA in complex with a deoxycholate ligand (DXC), as per 4ZP0.pdb (Heng et al., 2015).**

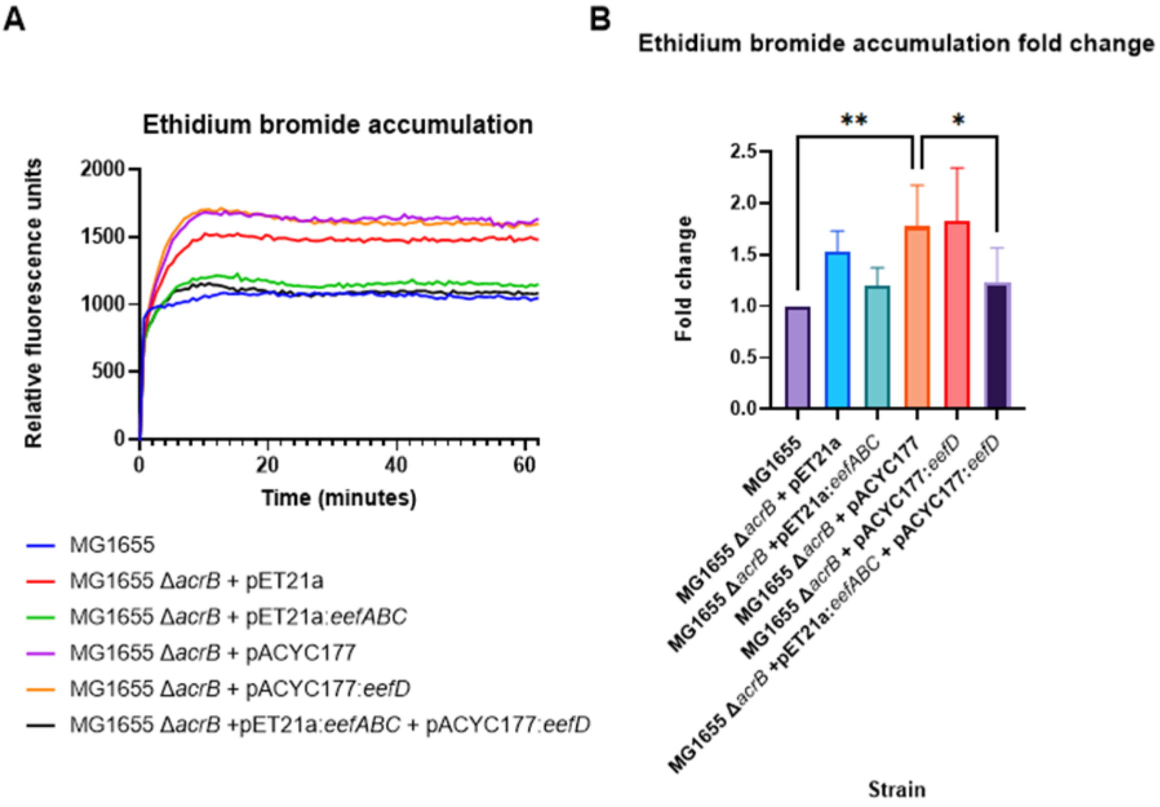


**Figure S16. Ethidium bromide accumulation assay for gain of function *E. coli* strains.** Panel A) Accumulation of ethidium bromide inside cells over time. Panel B) Fold-change in end point fluorescence value. Level of significance has been calculated using one-way-Anova analysis and compares the end point of all strains to the wildtype. Each data point is an average of 3 biological repeats, error bars show standard deviation.

**A**

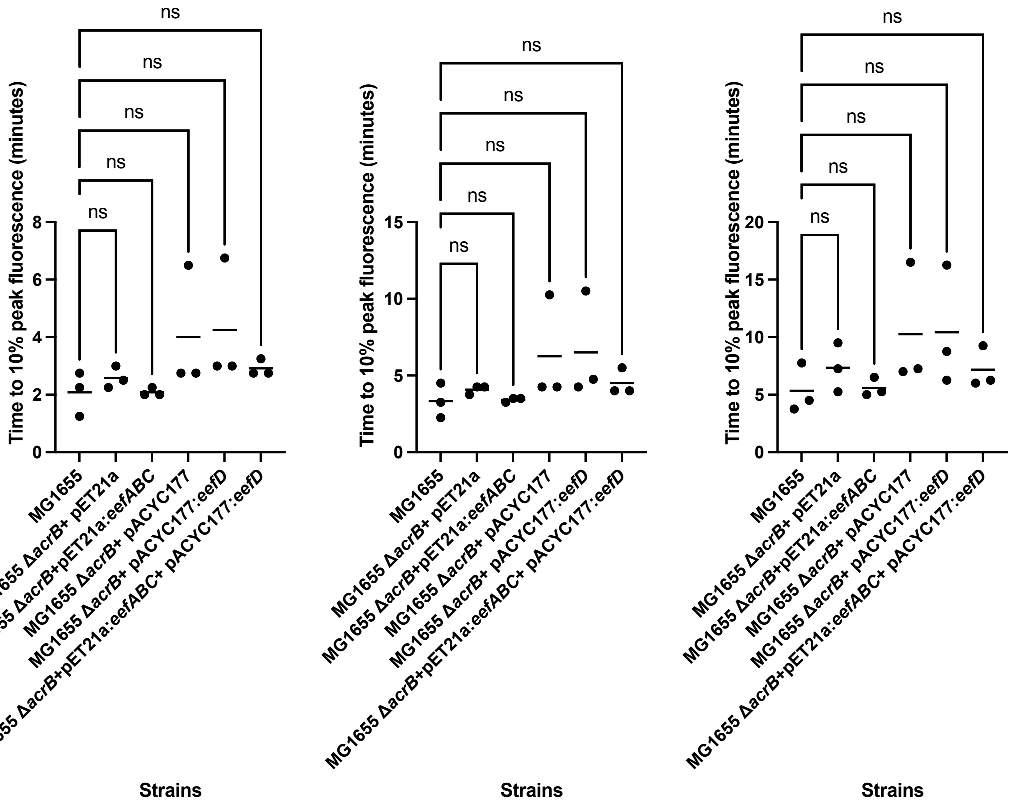


**B**

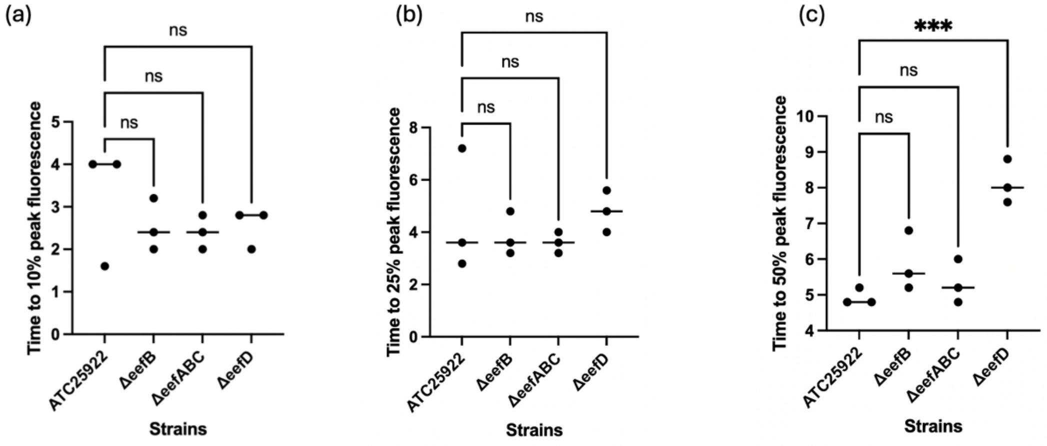


**Figure S17. Ethidium Bromide efflux for gain and loss of function *E. coli* strains.** The time taken for the volume of ethidium bromide inside the cell to decrease by 10%, 25% and 50% respectively in A) gain of function and B) loss of function strains. Level of significance was calculated using one-way-anova analysis. Data points are the average of 3 biological repeats, error bars of standard deviation.

**Structural analysis of the EefABC models.**

**EefC – structure:**

Our initial sequence analysis revealed several intriguing and unexpected features of the EefABC pump. We first focused on EefC which, as expected, was revealed to be a TolC paralogue, and is a member of the outer membrane factor (OMF) family (Paulsen et al., 1997). Within the *E. coli* genome there have been only 4 OMF family members identified so far – namely *tolC*, *mdtP* (*yjcP*), *mdtQ* (*yohH*) and *cusC* (*ylcB*)(Sulavik et al., 2001). Sequence alignment of EefC with the endogenous *E. coli* OMFs revealed that it is relatively distant to TolC branch, but closely aligned to the metal pump associated CusC and MdtP/MdtQ branch of the family (See Fig. S6). MAFFT multiple sequence alignment and the corresponding phylogenetic tree revealed that EefC is most (see Fig. S6 and S7). CusC is produced in a single operon with the RND-metal pump CusBA and is implicated in resistance against Cu(I)/Ag(I) ions generated under anaerobic conditions (Franke et al., 2003).

Comparison with other OMFs of known structure, including the *Pseudomonas* OMF channels OprM ((Akama et al., 2004) (1WP1.pdb) ; (Monlezun et al., 2015) (4Y1K.pdb ) ), OprN (5AZO.pdb; 5AZP.pdb) and OprJ (5AZS.pdb) (Yonehara et al., 2016) , as well as the Salmonella ST50 (Guan et al., 2015) (5BUN.pdb) and VceC (1YC9.pdb (Federici et al., 2005) confirmed that the closest relatives within the wider OMF family appear to be the *Pseudomonas* proteins OprM and OprJ (see Fig. S7), followed by the *CusC*, and correspondingly the former were used as structural templates for homology modelling (see Fig. S7 ).

From this structural alignment it is immediately notable, that there are very few gaps in the alignment between OprM/J and Eef (Fig. S7), which is in stark contrast to the TolC (Fig. **S8**). That allows for almost direct mapping of the aligned sequences of EefC onto OprM/J. The only notable exceptions are the marked shift in the position of the beta-2 strand of the first beta hairpin, resulting in significant shortening of the extracellular loop1, and secondly, the rather shortened C-terminus, which is significantly truncated, especially comparing to the TolC-branch of the OMF family (Fig. S8). Side by side comparison of EefC vs TolC allows to identify the key differences between the two channels (Fig. S9).

Of particular note is that the primary gating loop sequences (located on the tips of H3/H4 and H7/H8 helical hairpins in TolC) are not conserved amongst the TolC paralogues within *E. coli* (Fig. S8), which is consistent with their functioning with dedicated PAPs (and correspondingly transporter pairs) and as a result not being cross-reactive.

The comparison of these gating-loop regions in EefC (which correspond to the tips of the H3/H4 and H7/H8 helices in TolC) reveals that they are highly divergent, suggesting that their locking mechanism is markedly different in EefC from that observed in TolC, and hence EefC is unlikely to work with PAPs recognizing TolC "out of the box", but rather likely only interacting with its cognate PAP EefA.

This is particularly prominent on the tip of the H7/H8 hairpin, which harbours several key residues responsible for gating the TolC channel. While Y362 (TolC) is critically conserved across the family, the R367 (TolC), which links across to H4 binding to the conserved D153 (TolC; D206 in EefC) forming the so called "primary gate" thus sealing the TolC channel, (Andersen et al., 2002a, Bavro et al., 2008) is substituted in EefC by a small non-polar residue G412 making such interaction impossible.

Furthermore, the so-called "secondary gate", formed of the prominently conserved double aspartate ring (which in TolC if composed of D371/D374 from all three protomers (TolC 1EK9.pdb numbering)), and which forms the basis of ion-selectivity in TolC (Andersen et al., 2002b, Schulz and Kleinekathöfer, 2009) is fully absent and substituted by a small-hydroxylated residues instead – namely T416 and T419 (YLTVLTA), a feature unique to the EefC, as although D371 position is variable across the OMF family, the D374 position is remarkably conserved (even within the OprM/CusC clusters), and the only other exception are the rather deviant AatA proteins associated with dispersin secretion discussed above (VITYSSV). This changes the electrostatic properties of the channel dramatically, which is further exacerbated by the presence of the bulky L415 and L418, which provide a hydrophobic seal of the periplasmic end of the channel. The inner lining of the EefC tube is also markedly more hydrophilic, lined with a number of serine residues.

It is important to note that the residue corresponding to the EefC R422, is conserved across *Pseudomonas* OprJ/OprM/OprN clade, as well as in the CusC group of OMFs, while it is markedly different in TolC-branch of the family (including MdtP and MdtQ). In the former group of OMFs, its role seems to be coordination of the upper-ring of Asp-residues forming the "secondary gate" electrostatic seal, however in

EefC the residue corresponding to the second Asp is T419, and such an interaction is impossible.

These are not the only similarities with the OprM/N/J family however. The EefC exhibits long-N-terminal tail, which is some 30 residues longer than the corresponding region of TolC, but very much in line with the length seen in OprN/J. Furthermore, the N-terminus of EefC contains a cysteine residue, which isn't present at all in the TolC branch of the family, but which is conserved in the OprM/CusC (Monlezun et al., 2015), where it is a target to lipidation, usually taking the form of palmitoylation, which facilitates its insertion into the outer membrane. Indeed, as seen in Fig. S9 we used the OprM structure 4y1k.pdb (Monlezun et al., 2015) to model the palmitoylation of the C-terminal cysteine.

Moving to the outer-membrane-inserted β-barrel domain, in comparison to the TolC, EefC features markedly shorter extracellular loops, in particular the L2 loop, which occludes TolC opening (Vaccaro et al., 2008); suggesting more-open state of the EefC channel. Such OMP loops are also prominent targets of protective antibodies (Domínguez-Medina et al., 2020), and the non-protruding loops can serve to avoid antibody restriction and LPS occlusion.

Finally, there are significant differences in the organisation of the equatorial domain, which in EefC is composed of the residue ranges P31-R69 (N-terminal tail and alpha-helix H2); G240-P276 (the long loop connecting the hairpin-forming helices H4 and H6, which includes a short helix H5); plus a very short C-terminal tail (G450-K457) that follows the H8 helix. The corresponding regions in TolC are again markedly different, with the N-terminal section contributing to the equatorial domain being much shorter (S20-P37 full-length TolC numbering); the central section being also slightly shorter (G209-P232), while the C-terminal tail being dramatically longer at over 60 residues (G427-N493). The features of EefC equatorial domain match nearly perfectly the OprJ/OprN leaving virtually no gap in the alignment.

**<u>EefA – structure:</u>**

Our attention then turned to the PAP protein in the EefABC system, EefA which was also analysed in a similar way, revealing a sequence identity of 53.85% to the *Pseudomonas* MexA; but only 50.13% against the *E.coli* AcrA. Due to the availability of more complete full-length experimental templates AcrA homology models were created both using MexA (2v4d.pdb; (Symmons et al., 2009) ) and AcrA (5v5s.pdb;

(Wang et al., 2017)) as a template (utilising both I-TASSER and Swiss-model tools) (Fig. S10). While neither of the models got to the same level of confidence as EefC, the alignments with the known PAP structures are readily interpretable allowing unambiguous interpretation of the protein features.

Alignment of EefA and AcrA results in a direct amino-acid match with only 2 gaps in the alignment, one in the unstructured N-terminal tail, and another at position Q221 (EefA), which has a 4-residue-long deletion relative to AcrA. This region corresponds to the C-terminal end of alpha-helix 3 that is flanking the beta-barrel domain in PAPs, and it is also notable that shortening the same position is observed in MdtA/MdsA branch of the family, however it is not predicted to affect RND-binding (McNeil et al., 2019).

Despite overall similarity with MexA/AcrA, there are distinctive differences in the organisation of the EefA, notably in its α-hairpin domain (Fig. S11). While the RLS motif, which is speculated to be involved in PAP-OMF interaction (Kim et al., 2010, Song et al., 2014) appears to be preserved: R120/L124/S131/D136 (vs R128/L132/S139/D144 in *E.coli*), there are a number of significant changes in the adjacent residues, notably R123 (K131); V125 (L133); D134 (E142), which are predicted to produce steric clashes that would preclude its direct compatibility with TolC, and hence, such residues are likely to be playing a discriminatory role engaging with EefC.

Another unique feature is the predicted close proximity of C115 and C142 in the α-hairpin domain, the fold of which create a plausible condition for the formation of a Cys-Cys bridge. While the role of such a bridge is difficult to predict, it is very distinctive and not observed in any other known PAPs.

**EefB - structure:**

The RND-transporter component of the EefABC is the EefB, a large integral membrane protein which is predicted to form a functional trimer similar to other transporters in the family. Alignment of the EefB with other RND transporters of known function, including AcrB (*E. coli*); MtrD (*N. gonorrhoeae*); MexB (*P. aeruginosa*); CusB (*C. jejuni*) and AdeB (*A. baumannii*) revealed that they are highly similar, with CusB and MtrD being most divergent, while AcrB and MexB being equidistant to EefB (see Fig. S12).

This close similarity (against MexB 56.8% identity in 1038 residues overlap; Score: 2995.0; Gap frequency: 0.5%; against AcrB 57.0% identity in 1033 residues overlap; Score: 2997.0; Gap frequency: 0.7% - fewer gaps in MexB) allowed to generate high-fidelity homology model of EefB (I-TASSER C-score 0.89; range -5,2) using MexB as a template, which was chosen due to the fewer gaps in the alignment (0.7% vs 0.5% for AcrB) and slightly higher residue overlap with EefB (1038 vs 1033 residue overlap respectively).

Overall, the alignment of with MexB (and AcrB) produces very few gaps (the longest is 2 residues long) allowing for unequivocal attribution of secondary structure elements (Fig. S12). As can be seen from the side-by-side comparison of the EefB and AcrB, both present a virtually identical architecture (Fig. S13) and an estimated RMSD of around 1.8 Å over the C-alpha backbone, with only notable differences being the shortened loop connecting TM helices α16 and α17 in EefB (residues 498-507); and the slightly shorter C-terminal tail. Consistent with this, the critical proton-relay residues found in MexB D407, D408, K939 and T976 (Guan and Nakae, 2001)(corresponding to D407, D408, K940 and T978 of AcrB); (Su et al., 2006); (Takatsuka and Nikaido, 2006) are preserved in EefD (D408, D409, K935 and T972 respectively).

While a full analysis of the drug binding pockets and efflux ducts of EefB is beyond the remit of the current manuscript, the Access and Deep pockets (AP and DP) (Nakashima et al., 2013, Ramaswamy et al., 2018) conservation suggests a closer relation to the MexB/AcrB rather than to MexY type transporters, with notable preservation of the key hydrophobic residues in the so-called phenylalanine cluster [numbering as per MexB, with EefB numbering in brackets]: F136 (F137); F178 (F179); G179 (G180); F610 (F606); F615 (F611); F617 (F613 – switch loop); F628 (F624) (Ramaswamy et al., 2018).

**EefD – structure**:

The last member of the operon is EefD, and our analysis indicated that this is a secondary transporter that belongs to the 2.A.1.2 group of Transporter Classification Database (TCDB) which groups MFS (Major Facilitator Super-)family transporter, related to MDR-function including the Bcr/CflA, EmrD and MdfA branches (Reddy et al., 2012) (Quistgaard et al., 2016). This resulted in high-confidence homology models, revealing a classic 6+6 TM-helical arrangement which is closely related to the general topology observed in MdfA (Adler and Bibi, 2002) (Fig. S14).

Comparison of the homology model of EefD with the available structures of EmrD (Yin et al., 2006) and MdfA (Heng et al., 2015, Nagarathinam et al., 2018) further confirmed that the EefD is more closely related to the latter (Fig. S15), however there are significant differences in the substrate binding cavity.

Notably, the conserved acidic residue D34 (MdfA) involved in protonation and substrate coupling cycle of the transporter is preserved, while the second acidic residue E26(MdfA), is instead substituted by an aliphatic A16, suggesting a difference in the cycle. Furthermore, the residue Y127 (MdfA) which makes contact with D34 in the outwardly-open state of the transporter (Nagarathinam et al., 2018) is substituted by a positively charged R117, making such interaction impossible.

Additional analysis reveals significant conservation of the so-called motif A, and the critical residue E132 (D122 in EefD) that anchors TM4 to it, however the interactions that are anchoring the motif A to the linker helix 6-7 (notably MdfA residues R198 and D211) are not conserved. Finally there are a number of changes along the central substrate-binding cavity, suggesting significant divergence of potential substrates between EefD and MdfA (see table below for details). These discrepancies make predictions of the possible substrates of the EefD problematic, although some overlap with MdfA can be expected (Lewinson et al., 2003) e.g. chloramphenicol/thiamphenicol and lipophilic cations such as ethidium.

Overall, our analysis of the EefABC reveals unexpected similarities to the tripartite pumps OprM-MexAB and OprJ-MexCD in *Pseudomonas* (Masuda et al., 2000), rather than to any endogenous *E.coli* paralogues, which may suggest that the precursors of these pumps have been acquired via a lateral gene transfer event, similar to some other plasmid encoded efflux/secretion systems (e.g. the EAEC virulence plasmid pAA2) (Imuta et al., 2008) and correspondingly may play a role in virulence of the *E.coli*.
